# Hydrogels with Independently Controlled Adhesion Ligand Mobility and Viscoelasticity Increase Cell Adhesion and Spreading

**DOI:** 10.1101/2024.09.23.614501

**Authors:** Abolfazl Salehi Moghaddam, Katelyn Dunne, Wendy Breyer, Yingjie Wu, E. Thomas Pashuck

## Abstract

A primary objective in designing hydrogels for cell culture is recreating the cell-matrix interactions found within human tissues. Identifying the most important biomaterial features for these interactions is challenging because it is difficult to independently adjust variables such as matrix stiffness, stress relaxation, the mobility of adhesion ligands and the ability of these ligands to support cellular forces. In this work we designed a hydrogel platform consisting of interpenetrating polymer networks of covalently crosslinked poly(ethylene glycol) (PEG) and self-assembled peptide amphiphiles (PA). We can tailor the storage modulus of the hydrogel by altering the concentration and composition of each network, and we can tune the stress relaxation half-life through the non-covalent bonding in the PA network. Ligand mobility can be adjusted independently of the matrix mechanical properties by attaching the RGD cell adhesion ligand to either the covalent PEG network, the dynamic PA network, or both networks at once. Interestingly, our findings show that endothelial cell adhesion formation and spreading is maximized in soft, viscoelastic gels in which RGD adhesion ligands are present on both the covalent PEG and non-covalent PA networks. The dynamic nature of cell adhesion domains, coupled with their ability to exert substantial forces on the matrix, suggests that having different presentations of RGD ligands which are either mobile or are capable of withstanding significant forces are needed mimic different aspects of complex cell-matrix adhesions. By demonstrating how different presentations of RGD ligands affect cell behavior independently of viscoelastic properties, these results contribute to the rational design of hydrogels that facilitate desired cell-matrix interactions, with the potential of improving *in vitro* models and regenerative therapies.

## 1. Introduction

Hydrogels are essential in tissue modeling and regeneration because of their unique ability to mimic the physical and biochemical environments found in human tissues.^1,2^ These water-rich, crosslinked polymer networks can be designed to resemble the extracellular matrix (ECM) and are an ideal platform for studying cell behavior and tissue dynamics *in vitro*.^3,4^ Hydrogels offer the potential to support cell growth and differentiation, facilitating the development of functional tissue constructs that can be used for a variety of biomedical applications.^5,6^ Synthetic polymers are often utilized for these systems because they can be easily modified to tune important hydrogel properties, such as viscoelasticity and degradability.^7,8^ However, many polymers do not inherently support cell adhesion, and these networks are typically modified with cell adhesion molecules, such as the RGD peptide sequence, to support cell growth and viability within the matrices.^9,10^

The complexity of the extracellular matrix presents a significant challenge in designing artificial matrices and identifying which matrix features are most important for promoting physiological processes within engineered systems is a major goal of biomaterials research.^11,12^ For over 20 years it has been well-established that the stiffness of hydrogel matrices is a critical design factor influencing cell behavior.^13,14^ Human tissues exhibit complex viscoelastic properties, and in the past decade it has become widely appreciated that the ability of matrices to relax stresses is also a significant design consideration.^15,16^ Cells sense matrix properties through dynamic protein assemblies, such as focal adhesions, where integrins bind the local matrix and serve as mechanical links between the ECM and the cytoskeleton.^17,18^ The formation and maturation of cell-matrix adhesions is a complex and intricate process, characterized by the continuous movement of integrins in and out of integrin adhesion complexes.^19-21^ Integrins cluster during adhesion maturation,^22^ including the migration of ECM-bound integrins,^18,23^ and mechanical forces generated by the cells are a central unifying feature underlying nearly all aspects of cell adhesions (Figure 1 A-B).^16,24,25^ Many cell culture systems utilize cell adhesion ligands, like the RGD peptide, that are covalently attached to crosslinked polymer matrices.^26,27^ But studies show that the ability of cells to move adhesion sequences across the cell membrane is important for adhesions.^28-30^ Furthermore, recent research has highlighted that the degree of mobility of adhesion ligands is an important criteria for cell-matrix interactions.^31-33^

**Figure 1.**
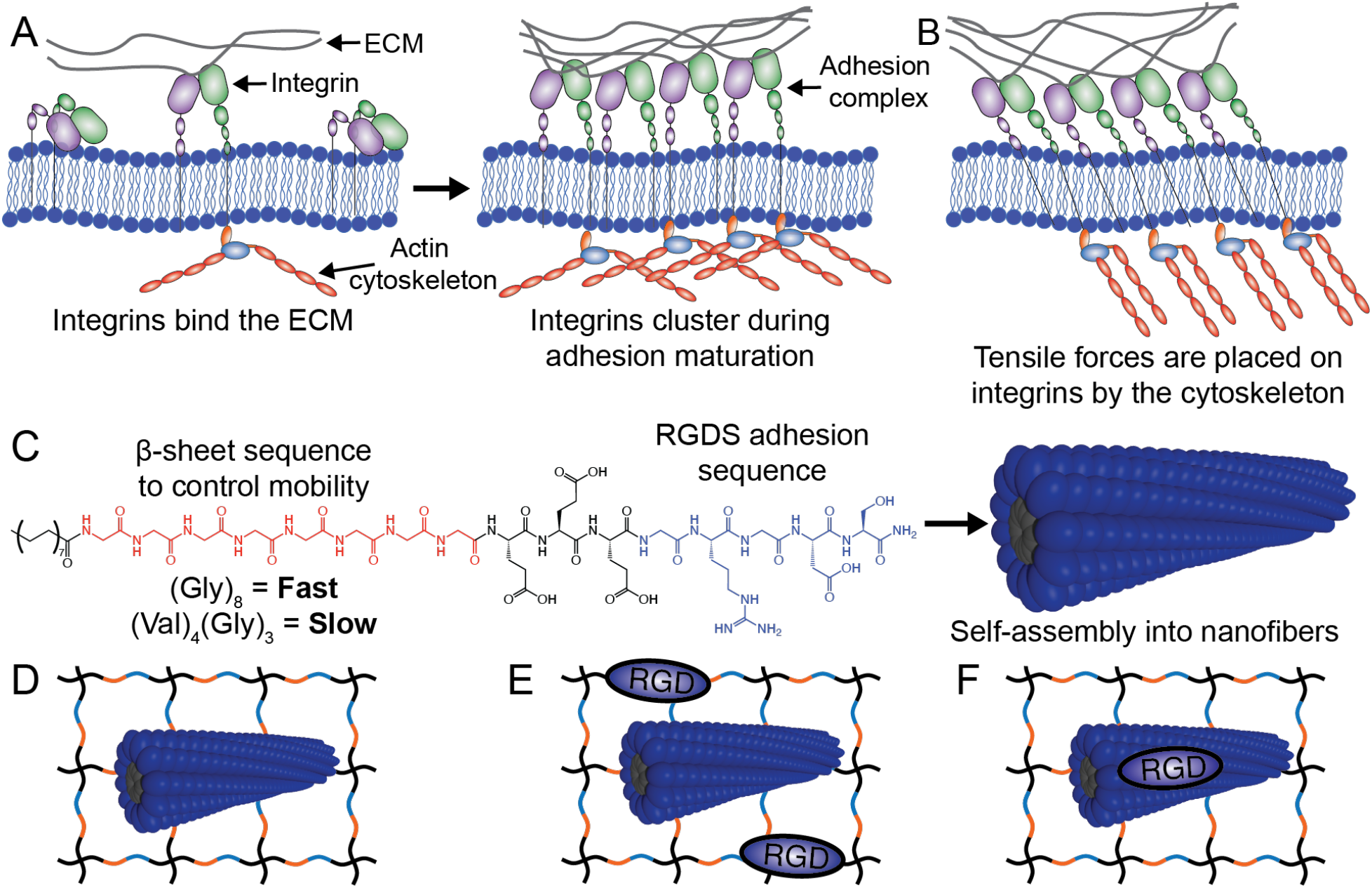
A) Cells adhere to the local matrix using integrins. Activated integrins bind the local matrix, which leads to integrin clustering and the formation of adhesion domains. B) Cytoskeletal forces are placed on the integrins and have a vital role adhesion growth. C) Peptide amphiphiles (PAs) are molecules that can self-assemble into nanofibers. They feature a hydrophobic alkyl tail, a β-sheet region which modulates intramolecular bonding, and can be functionalized to display cell adhesion ligands, such as the RGDS peptide. Modifying the β-sheet sequence can control the how dynamic peptides are within the nanofiber. D) PAs can be incorporated within a covalent PEG matrix to form an interpenetrating network with tailored viscoelastic properties. The mobility of RGD ligands can be tuned by coupling the adhesion ligands to E) the covalent PEG network or F) the dynamic PA network.

Although matrix stiffness, stress relaxation, and the mobility of adhesion ligands are recognized as crucial variables for cell-matrix interactions, our understanding of the relative importance of each factor is constrained by difficulties in isolating specific variables within biomaterials systems. Dynamic bonding, which can both enable the relaxation of stresses and enable cells to rearrange adhesion ligands, is typically engineered into synthetic matrices through either dynamic covalent bonds,^12^ the incorporation of non-covalent bonds, such as metal-ligand interactions in alginate,^15^ or self-assembling supramolecular polymers.^30,32^ These are frequently fabricated as single network matrices,^34^ and a challenge for these systems is that tuning the rate of dynamic bonding will generally modulate both the viscoelastic properties of the matrix and the mobility of adhesion ligands. Hydrogels can also be fabricated with multiple interpenetrating polymer networks (IPNs).^35,36^ An advantage of using these systems is that the properties of each network can be independently adjusted, enabling precise control over hydrogel features, such as the mechanical properties.^37,38^ IPNs can be made with both covalent and non-covalent supramolecular polymer networks, which is mostly commonly done by incorporating self-assembling peptides nanofibers into covalent polymer networks to form highly modular hydrogel platforms.^39-42^

This study introduces a synthetic hydrogel platform where we can independently tune storage modulus, stress relaxation, and the mobility of cell adhesion peptides. This is achieved by creating hydrogels composed of two interpenetrating polymer networks: a covalently crosslinked poly(ethylene glycol) (PEG) network and a supramolecular self-assembling peptide amphiphile (PA) network. By adjusting the proportions and components of each network, we can tailor the gel’s viscoelastic properties, including stiffness and stress relaxation. Attaching RGD adhesion ligands to either the covalent PEG or dynamic PA network allows for tuning the mobility of RGD ligands independently of the viscoelastic properties. Human umbilical vein endothelial cells (hUVECs) cultured in these hydrogels showed optimal spreading and adhesion in softer gels with rapid stress relaxation. Notably, focal adhesion formation and endothelial cell spreading were optimized in systems in which the RGD ligands were attached to both the dynamic self-assembled network and the covalent PEG network. Integrin-matrix interactions are complex, these findings highlight the need to recapitulate multiple aspects of cell-will help aid in the design of better biomaterials to control cell-matrix interactions.

## 2. Results and Discussion

### 2.1 Hydrogel Design and Characterization of Hybrid Hydrogel with Multiplexed Nanofibers

We designed a hybrid hydrogel featuring interpenetrating covalent and non-covalent polymer networks, allowing us to independently adjust hydrogel properties important for cell-matrix interactions, such as viscoelasticity and RGD ligand mobility. Poly(ethylene glycol) (PEG) is a synthetic polymer that is widely used within hydrogels^43^ and in clinical therapies,^44^ and PEG hydrogels are commonly fabricated by covalently crosslinking multi-arm PEG macromers with peptides that are cleaved by cell-secreted proteases.^45,46^ Peptide amphiphiles are a versatile platform for self-assembly.^47^ The feature a hydrophobic alkyl tail coupled a β-sheet-forming region that promotes the formation of high aspect ratio nanostructures, with β-sheets extending down the long axis of the nanofiber (Figure 1C).^48,49^ Mixing two different polymer networks has been shown to enable significant control over hydrogel properties,^35^ and our PEG-PA system has several variables that enabled the mechanical properties of the hydrogel to be modulated (Figure 1D). This includes the concentration of either the PA or PEG network, the peptide sequence within the peptide amphiphile, or the fraction of the PEG arms that were covalently crosslinked. 8-arm PEG macromers will typically form gels once more than 35% of the PEG arms are crosslinked with a peptide.^50^ The PA network features dynamic non-covalent bonding between the PA molecules, and this network stiffens the hydrogel but also enables the relaxation of stresses placed upon the matrix.

The presence of two discrete networks enables control the mobility of RGD cell adhesion ligands independently of the viscoelastic properties. Covalently attaching the RGD ligand to the PEG network renders it immobile (Figure 1E), while functionalizing the PA network with the adhesion ligand enables cells to dynamically arrange the peptide sequence during adhesion maturation (Figure 1F). The extent of RGD peptide mobility can be further controlled by altering the peptide sequence within the β-sheet region of the peptide amphiphile.^31-33^ In this work PAs with a highly mobile (Gly)_8_ β-sheet sequence (C_16_-G_8_E_3_GRGDS) are “Fast-RGD” peptides, and PAs with a lower mobility (Val)_4_(Ala)_3_ β-sheet sequence (C_16_-V_4_A_3_E_3_GRGDS) are “Slow-RGD” peptides (Figure 1C).

We used two different approaches to quantify the effect that the two different β-sheet sequences had on RGD mobility. In the first, fluorescence depolarization was performed by incorporating the hydrophobic fluorophore 1,6-diphenyl-1,3,5-hexatriene (DPH) into the hydrophobic core of the nanofiber. DPH emits fluorescence whose degree of polarization depends upon the local microviscosity, and more dynamic environments have lower viscosity which reduces the amount of fluorescence polarization. This was quantified using fluorescence depolarization and we found that the amount of DPH fluorescence anisotropy in the Fast-RGD PAs was approximately half the amount as found in the Slow-RGD PA, 0.18 to 0.36, respectively (Figure S1A). We also used transverse-relaxation nuclear magnetic resonance (T2-NMR) to measure the molecular mobility of molecules within the PA nanofibers. The relaxation rate of the protons on the methyl carbon of the palmitic acid hydrophobic tail and the methylene carbon on the aspartic acid side chain were analyzed and we found that the protons on the Fast-RGD had slower relaxation rates, which indicates that they have greater supramolecular motion (Figure S1B).

All hydrogels in this work were functionalized with 1.5 mM of a GRGDS cell adhesion ligand. The GRGDS peptide was either entirely on the PEG network (PEG-RGD), entirely on the PA network (Fast-RGD or Slow-RGD), or evenly split between the PEG and PA networks (Fast-RGD+PEG-RGD or Slow-RGD+PEG-RGD), with 0.75 mM on each. The total concentration of PAs was also constant across conditions, and the PA molecules were either 100% RGD-functionalized, 50% RGD-functionalized and 50% unfunctionalized, or 100% unfunctionalized with RGD. The degree of nanoscale RGD clustering is an important factor in determining how cells interact with their surrounding matrix.^22,51,52^ To ensure that the molecular density of the RGD sequence on the nanofibers was consistent across conditions, we designed the RGD-modified and unmodified PAs to prevent mixing at the molecular level. This was done by synthesizing RGD-modified peptides using amino acids having the L-stereoisomer and making the unfunctionalized PAs with amino acids having the D-stereoisomer (Figure 2A). The β-sheets that run down the long axis of PA nanofibers have a helical geometry.^48,49^ We postulated that peptides with opposite stereochemistries would sort into separate fibers at the molecular level, since the L-and D-PAs would form helices having opposing handedness.

**Figure 2.**
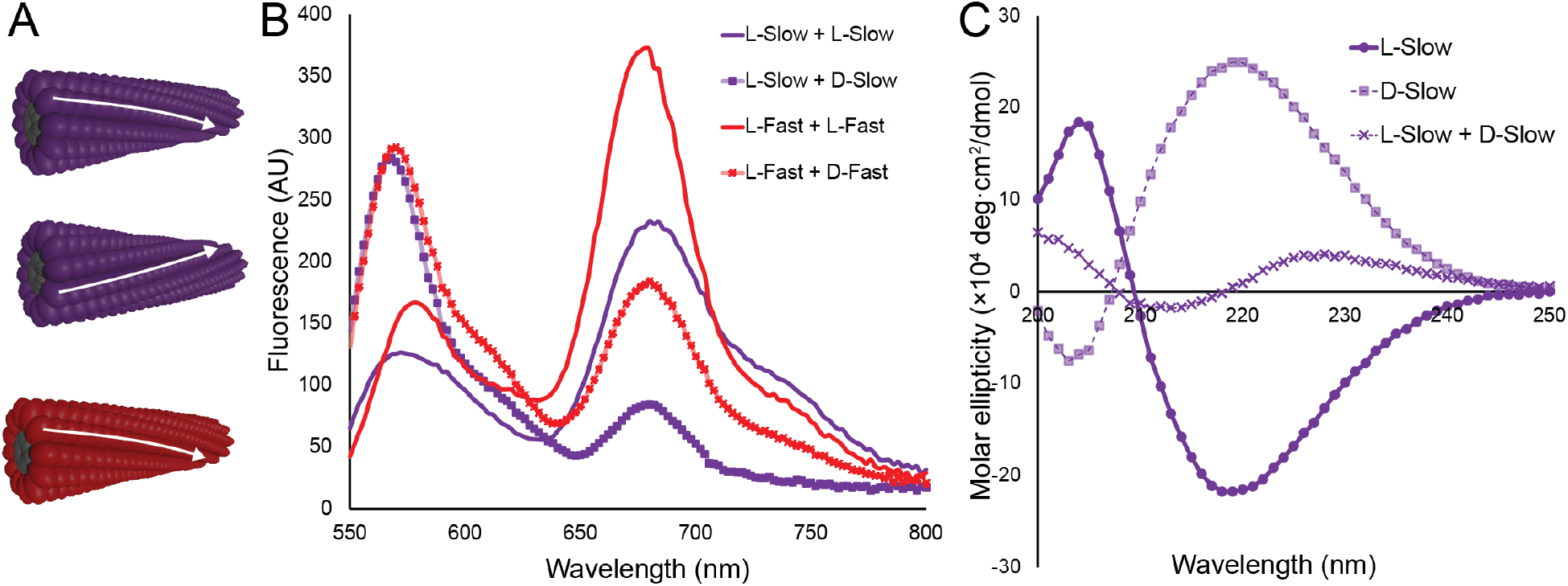
A) Orthogonal nanofiber networks can be formed by mixing L- and D-stereoisomers of peptide amphiphiles (purple fibers). The properties of PAs can be changed by changing the PA sequence (purple and red fibers) B) FRET PAs were synthesized with either a Cy3 or Cy5 fluorophore. Mixing L-versions of either the Fast or Slow PAs with Cy3 and Cy5 labelled molecules lead to FRET emission at 680 nm, while mixing the L- and D-versions had significantly reduced emission, indicating reduced mixing at the molecular level. All samples were excited with 515 nm light. C) Circular dichroism shows that the L- and D-versions of the Slow PA form β-sheets with opposite chirality.

We quantified the amount of molecular mixing at the nanofiber level by synthesizing L- and D-versions of both the Fast and Slow PAs that were labelled with either the Cy3 or Cy5 fluorophores. Cy3 and Cy5 will undergo Förster resonance energy transfer (FRET) when in close proximity to each other, with the pair having a Förster radius (R_0_) of Cy3/Cy5 is 52Å.^53^ When we attached the Cy3 and Cy5 flurophores to peptides having the same L-stereochemistry we see a Cy5 emission peak around 680 nm that is significantly larger than the Cy3 emission at 575 nm for both the Fast and Slow PA molecules (Figure 2B). This data indicates that there is significant mixing at the molecular level within the nanofibers, which is in line with previous studies.^54^ However, mixing L- and D-versions of the same peptide sequence leads to a diminished FRET signal, with the Cy3 emission at 575 nm being significantly greater than the Cy5 emission at 680nm when excited with 515 nm light. These results show that there is less mixing on the molecular level, likely due to the presence of mostly homogenous nanofiber networks. While some FRET is still seen in the mixed systems, it should be noted that the nanofibers themselves in close enough proximity that a fraction of the fluorphores on different nanofibers are likely within the Förster radius of the FRET pair.

We also performed circular dichroism (CD) on the L- and D-versions of PAs, and in mixed systems with both stereoisomers present (Figures 2C and S2). The CD spectra from the Slow-PA, which has a V_4_A_3_ β-sheet sequence, shows that both the L- and D-versions form a canonical β-sheet signal, with the L-stereoisomer having a CD minimum at 219 nm and a maximum at 204 nm. The CD spectra from the D-stereoisomer is inverted, with a maximum at 219 nm and a minimum at 203 nm, validating that the β-sheets have the opposite twist. Mixing both the L- and the D-stereoisomers shows an attenuated signal, likely due to the different nanofibers cancelling out the absorption of circularly polarized light. The Fast-PA has a β-sheet sequence containing eight repeats of glycine, an amino acid which lacks a chiral center and has no CD signal. Interestingly, while the glutamic acid residues were either L-for the L-stereoisomer and D-for the D-stereoisomer, the CD signal for the different stereoisomers was shifted but not inverted (Figure S2).

### 2.2 Tuning the Stiffness and Stress Relaxation of the Hybrid Hydrogel

An advantage of the PEG-PA system is that there are multiple variables which can be tuned to adjust different properties of the hydrogel system (**Table 1**). The hydrogels in these studies were between 2%-4% 20 kDa 8-arm PEG by weight and had between 40%-95% of the PEG arms crosslinked with a GPQGIWGQ MMP-degradable peptide.^55^ Hydrogel stiffness was primarily influenced by the fraction of PEG arms crosslinked and the weight fraction of the hydrogel comprised of PEG, and increasing either lead to stiffer hydrogels (Figure 3). Endothelial network formation is optimized on very soft gels,^50^ and the gels in this work had storage moduli in the range of 160 ± 24 Pa to 246 ± 95 Pa, with the exception of the “Stiff” gel, which had a storage modulus of 667 ± 11 Pa (Figure 3A). The ability to quickly relax stresses was only observed in hydrogels with 40% of the PEG arms crosslinked by the MMP-degradable crosslinking peptide. Work across several different biomaterials systems suggests the stress relaxation half-life of hydrogel matrices is an important matrix parameter for controlling cell phenotype.^15,16,56^ In our studies the Fast-RGD and Slow-RGD systems relaxed half of their stresses by after 76 seconds and 208 seconds, respectively (Figure 3B). After 1,800 seconds (30 minutes) the Long-relaxation Gel still had 66% of the original stress, while the Stiff gel had 91%. Recent work with endothelial cells shows that hydrogels with that relax half their stresses in under 500 seconds had greater network formation than those that relax stresses in greater than 500 seconds.^56^ Interestingly, this is also seen with other cell types, as a stress relaxation half-life of 500 seconds seems to be a critical value for modulating the differentiation of mesenchymal stem cells.^57^ In our platform the Long-relaxation Gel had a storage modulus of 160 ± 24 Pa and has similar stiffness as the Slow-RGD and Fast-RGD, but with a more limited ability to relax stresses.

**Table 1.** Hydrogel compositions for the four viscoelastic conditions used within this work.

|  | PEG concentration | Fraction crosslinked | PA |
| --- | --- | --- | --- |
| Fast-RGD | 2.75% | 40% | Fast |
| Slow-RGD | 2.75% | 40% | Slow |
| Long-relaxation Gel | 2% | 95% | Fast |
| Stiff Gel | 4% | 90% | Fast |

**Figure 3.**
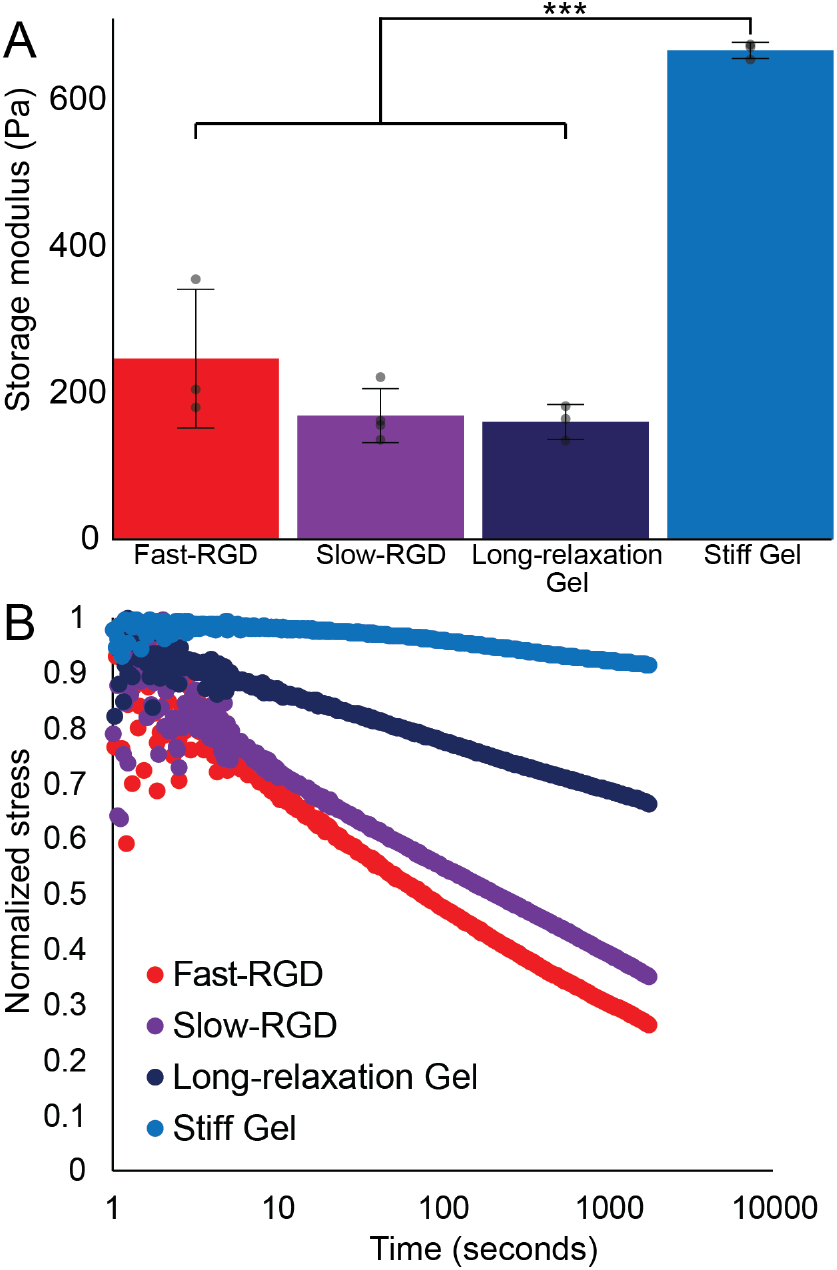
A) Increasing the concentration of PEG in the hydrogel leads to a stiffer gel having a storage modulus of 667 ± 11 Pa, versus 160-250 PA for the other softer gels. B) The stress relaxation half-life of the PEG-PA gels can tuned, with slow relaxing gels having a relaxation half-life in excess of 1,800 seconds, while the Fast-RGD and Slow-RGD relax half of the stresses in under 210 seconds.

### 2.3 Focal Adhesion Formation and Endothelial Cell Spreading in Hybrid Hydrogels

Endothelial network formation is generally maximized on very soft hydrogels^50^ and that matrices which both relax stresses and enable integrin clustering promote vascular morphogenesis.^29,56^ We cultured hUVECs in a series of PEG-PA hydrogels and either tuned the mechanical properties while keeping the adhesion environment the same, or modulated the adhesion environment while maintaining similar mechanical properties. To better understand how hydrogel properties influenced cell-matrix interactions, we stained the hydrogels after 24 hours of culture and imaged them using confocal microscopy (Figures 4 and S3). Vinculin, a protein found within focal adhesions, had the highest intensity per cell in hydrogels that have RGD ligands on both Fast-PA and PEG polymer networks (Fast-RGD+PEG-RGD). Hydrogels with Slow-RGD and PEG-RGD had slightly lower vinculin staining intensity comparable to the Fast-RGD+PEG-RGD gels, with approximately 95% of the staining intensity per cell (Figure 4U). Having RGD only on the Fast-PA network reduced vinculin staining per cell compared to the Fast-RGD+PEG-RGD, although the results were not significant (p = 0.31). Notably, both the Slow-RGD and PEG-RGD had significantly reduced vinculin staining per cell, with intensities only 49% and 55% of that of Fast-RGD+PEG-RGD, respectively.

**Figure 4.**
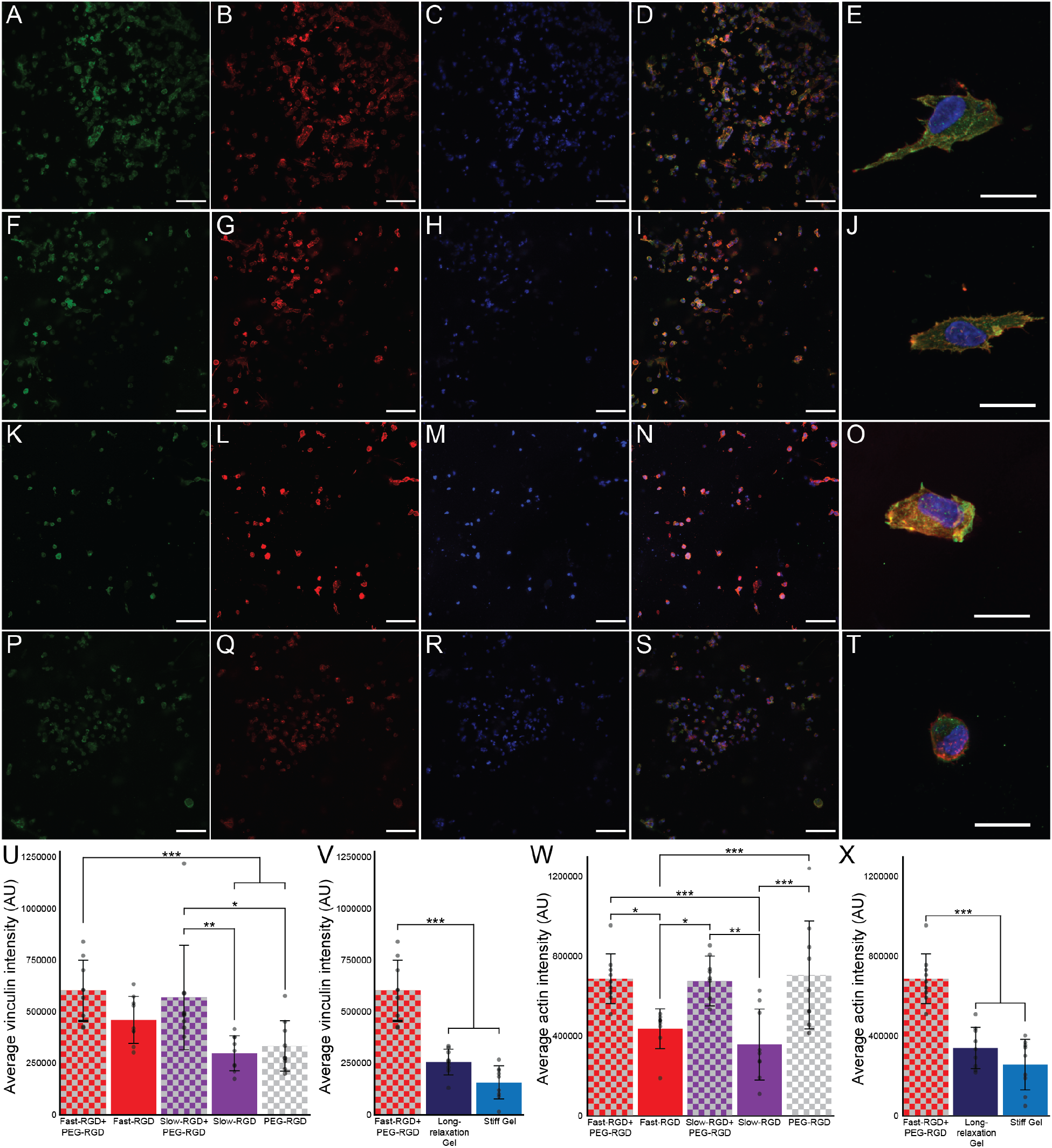
Immunocytochemistry was done on hUVECs within hydrogels after 24 hours of culture, staining for vinculin (green), a protein found in focal adhesions, in addition to actin (red) and the nuclei (blue). Fast-RGD + PEG-RGD (A-E), Slow-RGD + PEG-RGD (F-J), PEG-RGD (K-N), and Fast-RGD (P-T) were imaged. The vinculin intensity per cell was quantified as a function of (U) RGD presentation or (V) hydrogel viscoelasticity. Actin intensity was also quantified as a function of (W) RGD presentation or (X) hydrogel viscoelasticity. Scale bars are 100 µm, except for the right column (E, J, O, T), where they are 20 µm. Statistics are calculated from Tukey’s post-hoc test, with an N = 9, where * indicates p <0.05, ** indicates p<0.01, and *** indicates p<0.001.

We then kept the Fast-RGD+PEG-RGD adhesion system but changed the mechanical properties, and found that Slow Relaxing Gel, which had a similar stiffness as the Fast-RGD+PEG-RGD, but slower stress relaxation, had significantly decreased vinculin staining per cell, with only 42% of the vinculin intensity (p <0.001) of the Fast-RGD+PEG-RGD condition (Figure 4V). Increasing the stiffness of the gels further reduced vinculin staining, with only 25% of the intensity as soft, stress-relaxing gels.

The actin cytoskeleton was stained with rhodamine phalloidin, and actin intensity per cell was quantified after 24 hours. Interestingly, the condition with the most intense actin staining was the PEG-RGD condition in which all of the RGD peptides are covalently bound to the PEG hydrogel. The two other conditions in which RGD was attached to the PEG, Fast-RGD+PEG-RGD and Slow-RGD+PEG-RGD, had similar staining intensities per cell, within 5% of the PEG-RGD values. Fast-RGD and Slow-RGD, in which all RGD peptides are attached to the non-covalent PA network, had actin intensity that were 62% and 50% of the PEG-RGD, respectively. The long-relaxation gels and stiff gels had the lowest actin staining intensities, with the stiff being only 36% of the PEG-RGD.

Comparing the vinculin and actin staining intensity, an interesting finding is that the PEG-RGD had less vinculin staining per cell than the Fast-RGD condition, with only 72% of the staining, although the results were not significant (p = 0.48). However, PEG-RGD had significantly more actin staining than Fast-RGD (p = 0.01). The vinculin results suggest that having highly dynamic RGD ligands increasing focal adhesion formation, as measured by vinculin staining, while the actin network that cells use to generate stresses is maximized when the RGD ligands are covalently bound to the matrix. It is also apparent that adhesion formation and actin network structure depends upon both the presentation of RGD ligands and the viscoelastic properties of the hydrogel matrix.

hUVEC spreading was imaged and quantified on Days 1 and 7 using confocal microscopy (Figure S4 and 5). Endothelial cells in the soft, stress-relaxing hydrogels with both the Fast-RGD and PEG-RGD had the most area after 24 hours, and gels that had PEG-RGD had increased spreading compared to gels that did not (Figure S4). Increasing the stress relaxation half-life of the hydrogel decreased the cell area, and increasing hydrogel stiffness decreased it further. Cell spreading increased by Day 7, and cells in the Fast-RGD+PEG-RGD condition had significantly more spreading than any other condition (Figures 5 and S4). Slow-RGD+PEG-RGD had the second most cell spreading, indicating the importance of both dynamic and covalent RGD for maximizing endothelial cell area.

**Figure 5.**
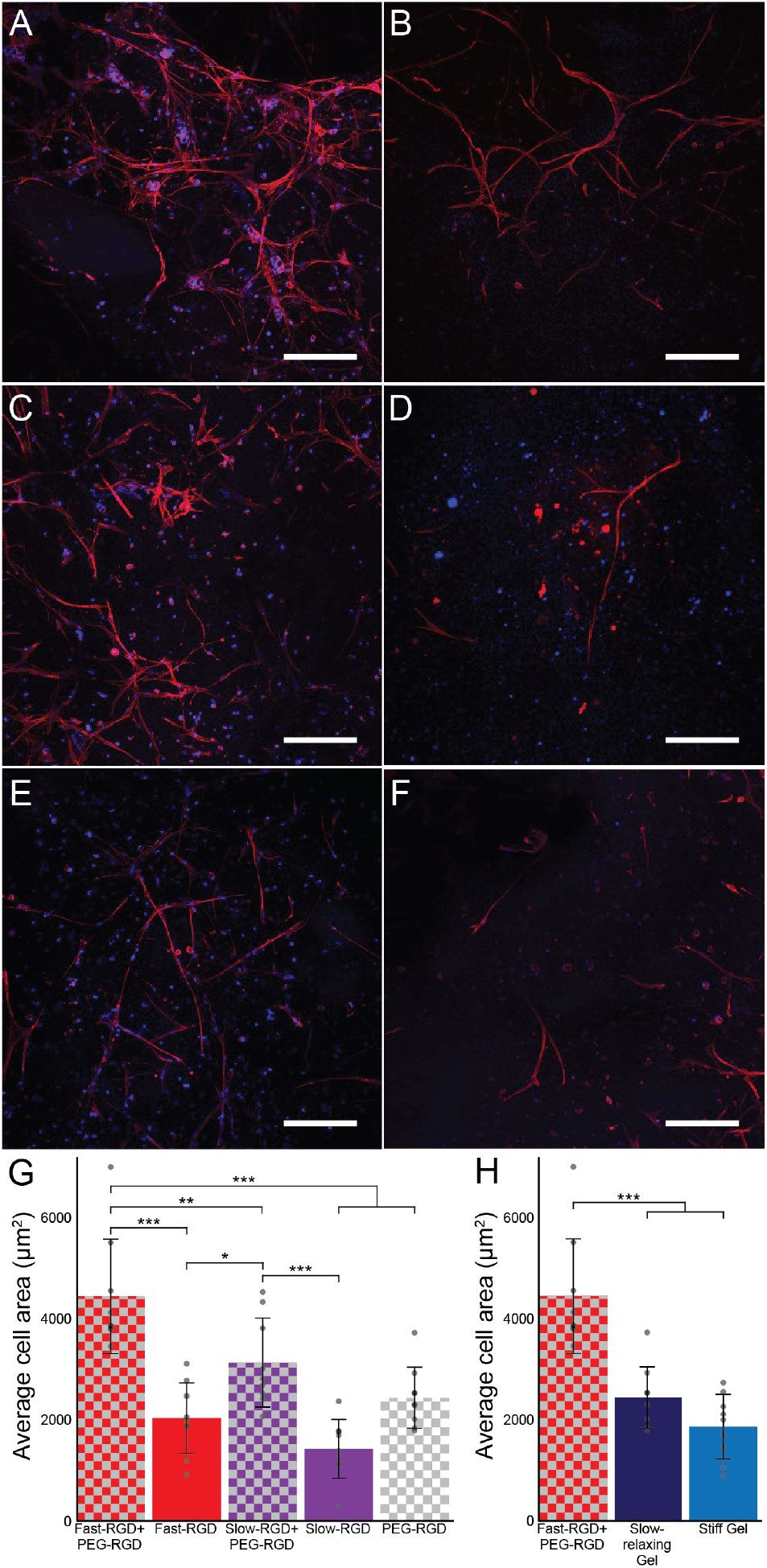
Immunocytochemistry was done on hUVECs within hydrogels after seven days of culture, staining for actin (red) and the nuclei (blue). A) Fast-RGD + PEG-RGD), B) Slow-RGD + PEG-RGD, C) PEG-RGD, D) Fast-RGD, E) Long-relaxation gels, and F) Stiff Gels were imaged. The average cell area was quantified as a function of (G) RGD presentation or (H) hydrogel viscoelasticity. Scale bars are 200 µm. Statistics are calculated from Tukey’s post-hoc test, with an N = 9, where * indicates p <0.05, ** indicates p<0.01, and *** indicated p<0.001.

The significant improvement in cell spreading observed in hydrogels containing both mobile and PEG-conjugated RGD highlights the complexity of cell adhesion. Adhesion formation is a dynamic process where ligand-bound integrins migrate into focal adhesions, applying forces to the local matrix that strengthen and grow these assemblies.^18^ Integrin-fibronectin bonds can sustain forces of approximately 93 pN before rupture,^58^ while the covalent C-O and C-C bonds found in PEG can support forces exceeding 2, 000 p N.^59^ Therefore, the forces placed on integrins will almost certainly disassociate the non-covalent integrin-matrix interactions before breaking the covalent bonds in the polymer network. However, self-assembled peptide nanofibers are highly dynamic and both previous work,^54,60^ and our own results in Fig. 2B indicate that individual PA molecules frequently exit one nanofiber and insert into a different nanofiber, as indicated by the FRET signal when homogenous fibers are mixed. The ability of cells to rearrange integrin-ligand complexes in covalent PEG-RGD is likely to be limited to local deformation of the polymer networks, while PA molecules can move throughout the gel. On the other hand, highly dynamic networks may not be able to support the significant cell-mediated forces that are important adhesion maturation.^61^ In addition to the inherent dynamicity of self-assembled systems, cells frequently apply strains of 3-4% and up to 20-30% on their local matrix during adhesion and migration,^15,62,63^ while many self-assembled hydrogels undergo mechanical failure above 1-2% strain.^49^ As a result, the forces cells can place on the local matrix is likely limited by the weak covalent bonding in the RGD-PA networks, whereas in the RGD-PEG network it is limited by the integrin-RGD interactions.

Having multiple networks functionalized with integrin-binding peptides may be advantageous because each network can be optimized for a specific aspect of cell-matrix adhesions. Dynamic networks can support integrin clustering and focal adhesion formation, while covalent networks are likely able to support much larger forces, and most covalent hydrogels can undergo significant strains without failure.^64^ It is also notable that studies that have tuned adhesion ligand mobility find that increasing mobility increases the formation of adhesions.^31,32^ Molecular mobility is typically enhanced by either increasing the rate of bond exchange or using weaker bonds between molecules, both of which are likely to reduce the ability of the ligand to sustain applied forces. The need for highly mobile ligands that can withstand significant forces is a difficult design challenge for single-component systems, and multi-component biomaterial systems that can mimic multiple aspects of cell-matrix interactions are likely necessary to replicate the complex and dynamic interactions cells have with their surroundings.

## 3. Conclusion

In conclusion, we developed a hydrogel platform consisting of a covalent hydrogel network and interpenetrating self-assembled peptide networks. We were able to tune the viscoelastic properties, including both hydrogel stiffness and stress relaxation, by altering the concentration of the PEG network and the extent of covalent crosslinking within the PEG network. We found that peptide amphiphile nanofibers with amino acids having either the L-or D-stereochemistry enabled the formation of discrete nanofiber networks which had reduced mixing at the molecular level. These PEG-PA hybrid hydrogels were functionalized with RGD adhesion ligands, and the mobility of the ligands was tuned independently of the mechanical properties. We found that endothelial cell spreading was maximized in soft hydrogels that quickly relaxed stresses. We also found that within soft, stress-relaxing hydrogels, endothelial cell spreading and adhesion formation was optimized in gels in with the RGD cell adhesion ligand was present on both the covalent PEG network and dynamic PA network. Furthermore, having highly mobile PA molecules was advantageous than PAs with lower molecular mobility.

In summary, this study reports the design of a highly modular hydrogel system that can be used to understand how specific matrix features influences cell-material interactions within engineered systems. The data showing that hydrogels dynamic and static RGD adhesion ligands have improved biological properties is a key finding that highlights that systems with only a single presentation of the RGD ligand may be unable to fully recapitulate the different aspects of adhesion maturation. Since self-assembled peptide networks can be easily integrated into any covalent hydrogel matrix, this work has the potential to improve the physiological relevance of many in vitro cell culture platforms.

## Acknowledgements

We would like to acknowledge our funding sources, the NIH (1R21GM143593-01) and NSF (Award 2138723). We are grateful to the lab of Lesley Chow for the use of their preparative high performance liquid chromatography, and to Samuel J. Rozans for synthesizing the Cy3 and Cy5 fluorophores.

## Supporting Information

Detailed methods, additional data, chemical characterization, and statistical analyses can be found in supporting information.

## Notes

### Competing Interest Statement

The authors have declared no competing interest.

