## Supplemental Information for "Hydrogels with Independently Controlled Adhesion Ligand Mobility and Viscoelasticity Increase Cell Adhesion and Spreading"

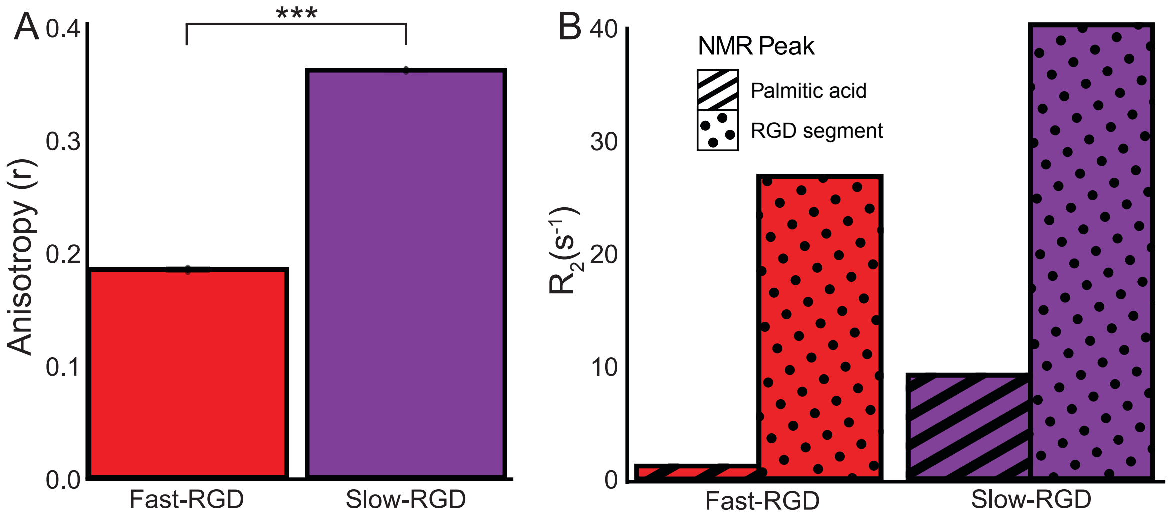


**Figure S1.** (A) Fluorescence depolarization of the 1,6-diphenyl-1,3,5-hexatriene (DPH) within Fast-RGD and Slow-RGD nanofibers shows a lower degree of anisotropy within Fast-RGD peptide amphiphile nanofibers compared to Slow-RGD peptide amphiphile nanofibers. (B) The Fast-RGD had a slower relaxation rate in T2-NMR. This indicates a higher degree of molecular mobility within Fast-RGD peptide amphiphile nanofibers.

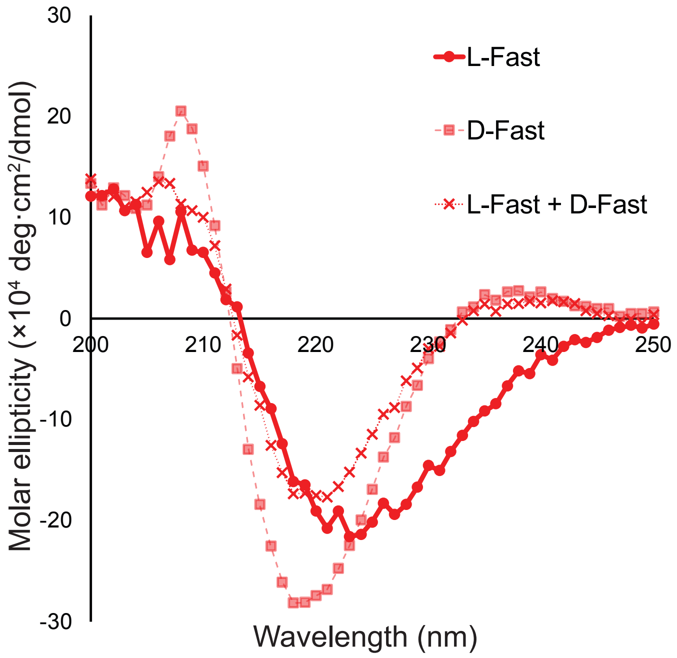


**Figures S2.** Circular dichroism on the L- and D- version of the C16-G8E3 “Fast” PA.

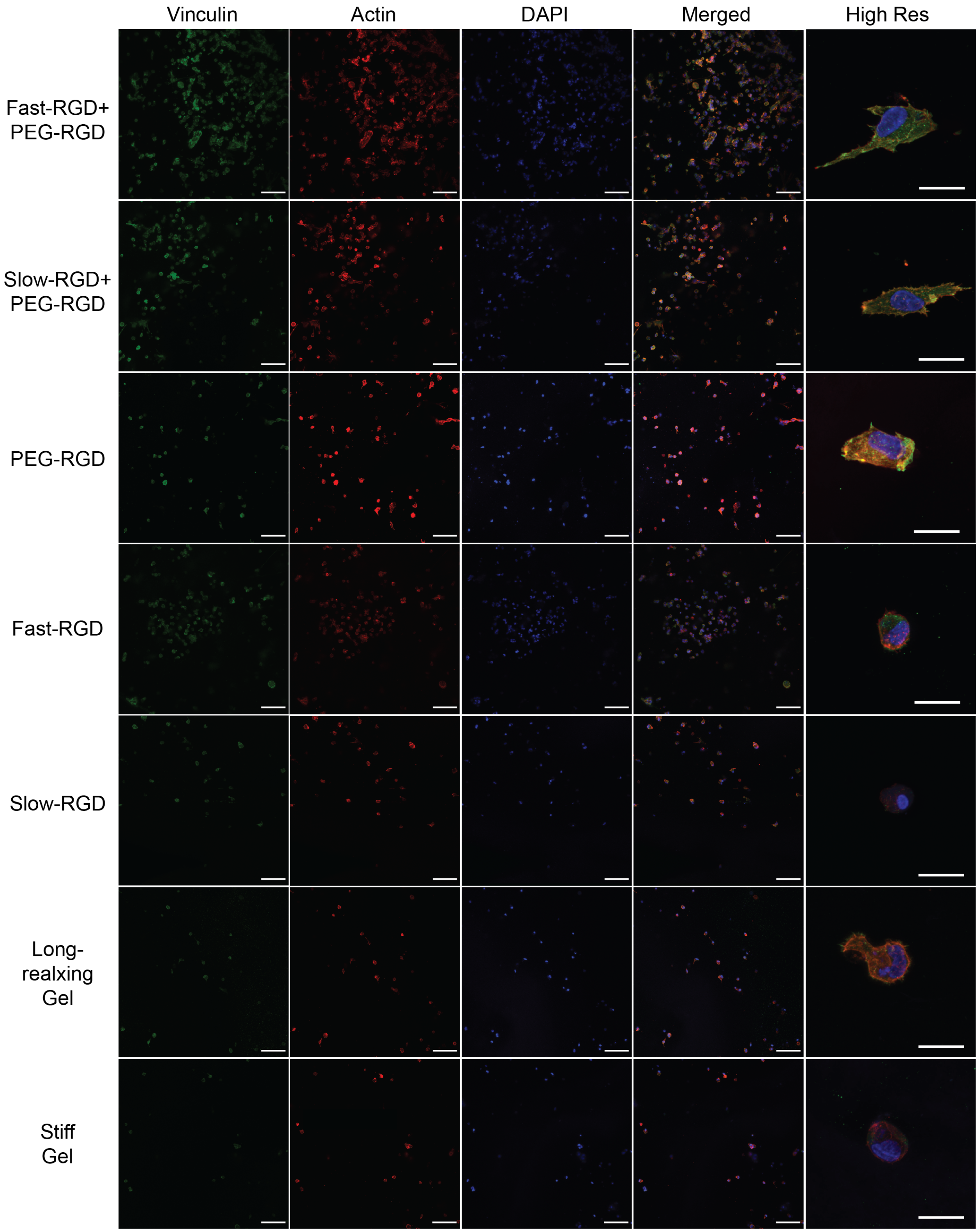


**Figure S3.** Immunocytochemistry of hUVECs at 24 hours staining for vinculin (green), actin (red) and DAPI (blue). The left four panels are low magnification (scale bar is 100 μm), the right panel is higher magnification (scale bar is 20 μm).

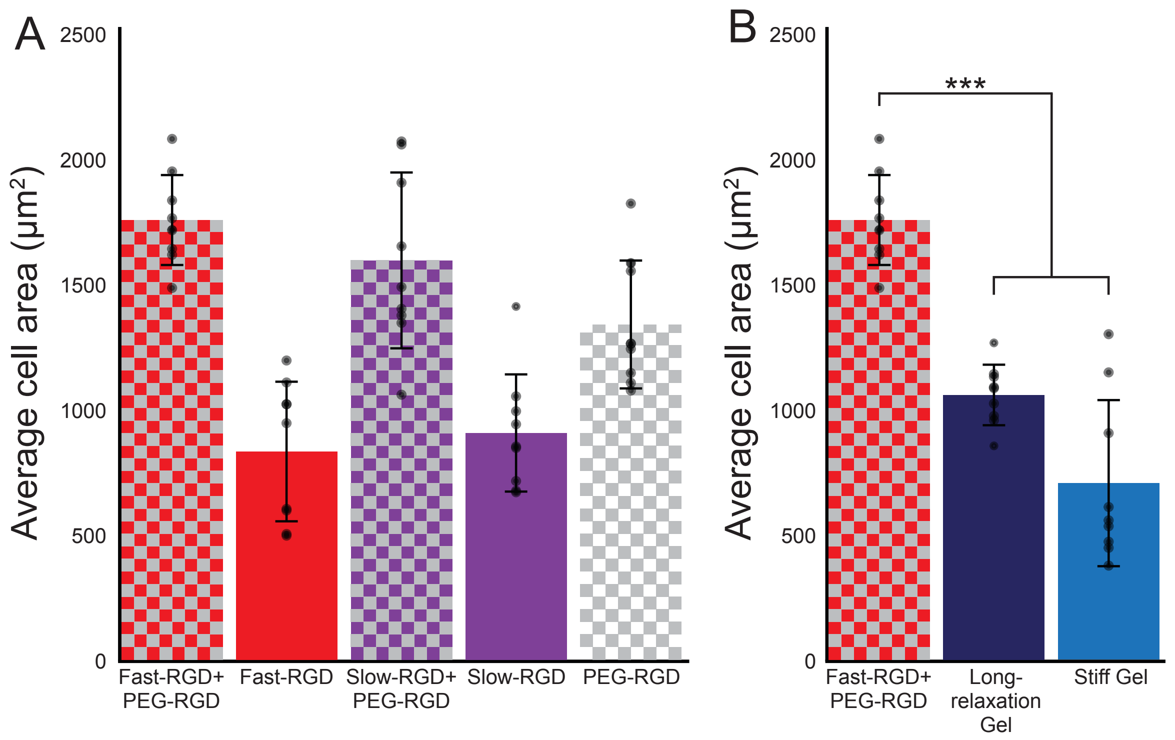


**Figure S4.** Average hUVEC area after 24 hours of culture within hydrogels for different RGD presentations (A), or hydrogels with different viscoelasticity (B). *** indicated $p<0.001$ by a

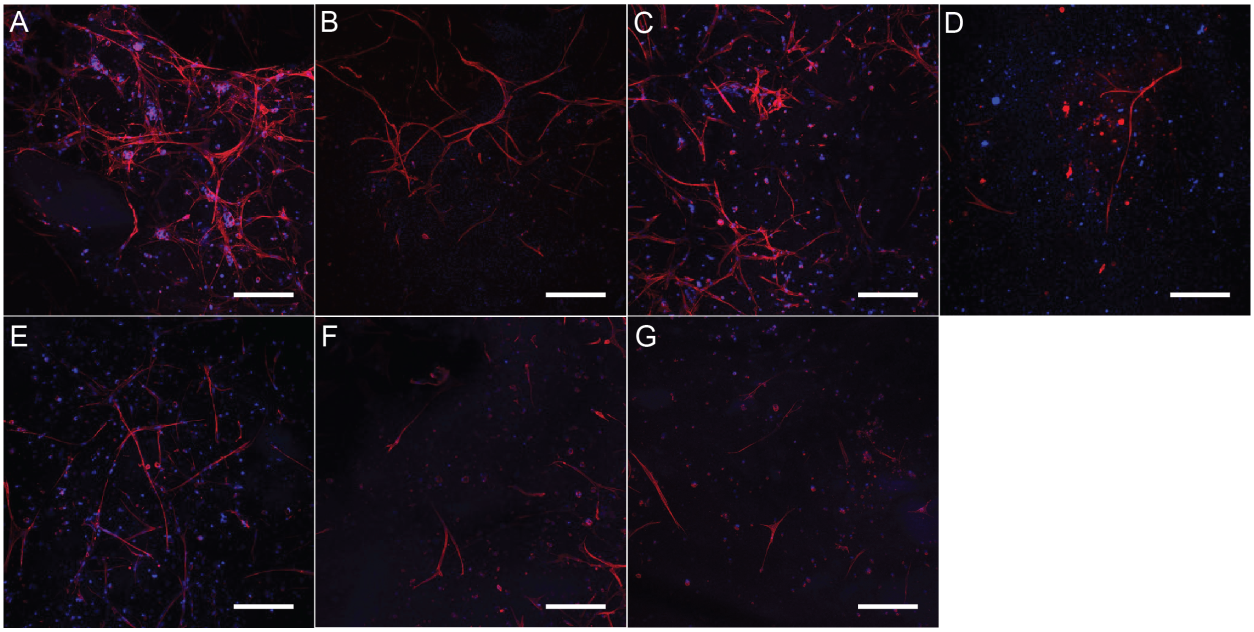


**Figure S5.** Immunocytochemistry of hUVECs at seven days staining for actin (red) and DAPI (blue). A) Fast-RGD + PEG-RGD), B) Slow-RGD + PEG-RGD, C) PEG-RGD, D) Fast-RGD, E) Long-relaxation gels, F) Stiff Gels, and G) Slow RGD (scale bar is 200 μm).

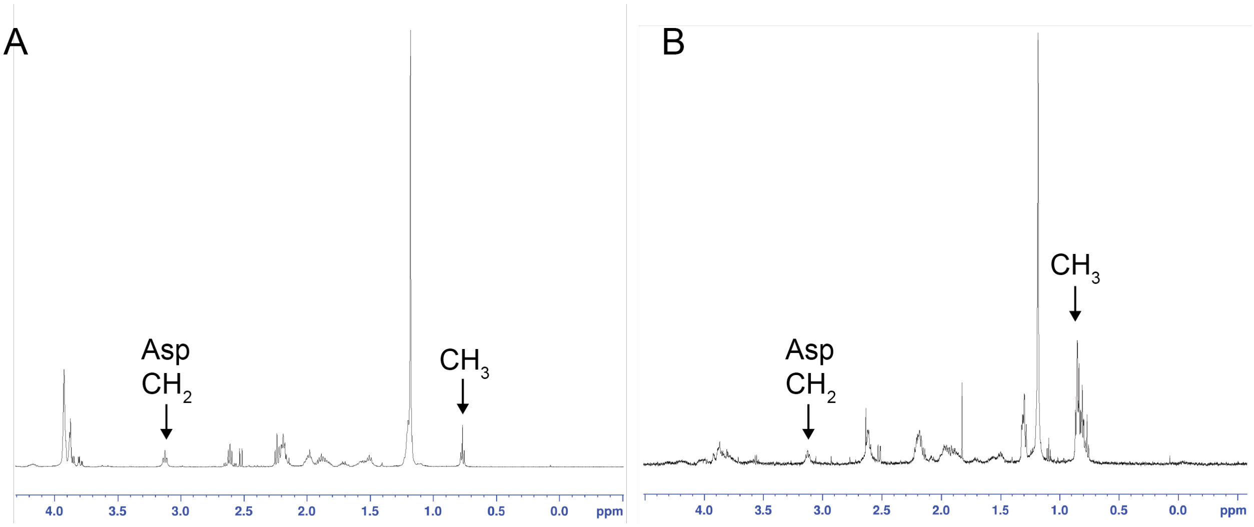


**Figure S6.** NMR spectra for A) Fast-RGD PA and B) Slow-RGD PA, with peak assignments for the methyl group on the palmitic acid tail, and methylene group found within aspartic acid.

### <u>Materials and methods</u>

#### Materials

All peptide synthesis reagents were purchased from Chemscene or Ambeed. N,N-Dimethylformamide (DMF) and dichloromethane (DCM) were obtained from VWR BDH Chemicals, while piperidine and trifluoroacetic acid were supplied by Millipore Sigma. Diethyl ether was acquired from Fisher Scientific, and N,N-Diisopropylethylamine (DIPEA) from VWR. The 20 kDa 8-arm poly(ethylene glycol) dibenzocyclooctyne (PEG-DBCO) was purchased from Biopharm PEG. To purify it, the PEG was dissolved in isopropanol and impurities were removed using a 10 kDa Amicon centrifugal filter. Specifically, approximately 200 mg of PEG was dissolved in 5 mL of isopropanol and centrifuged at 4,500 RPM until the volume of PEG-DBCO/ isopropanol in the filter was reduced to less than one milliliter. This process was repeated twice. The resulting filtrate was lyophilized and subsequently used in the experiments.

#### Methods

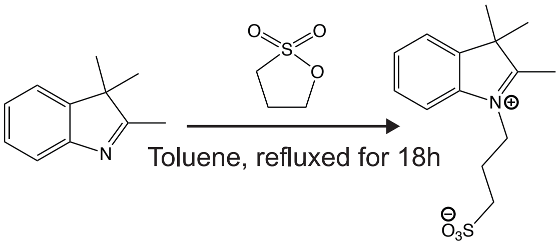


*Cyanine Dye Synthesis*

*Synthesis of sulfonated indolenine:* A mixture of 2,3,3-trimethylindolenine (20.0 mL, 127.0 mmol) and 1,3-propanesultone (15.5 g, 127.0 mmol) was refluxed in toluene (500 mL) for 18 h, which resulted in the formation of a dark red precipitate. The reaction was then cooled to room temperature and was concentrated under reduced pressure. The product was then dissolved in dichloromethane and then precipitated in ethyl ether in 50 mL centrifugal tubes. These were then centrifuged to concentrate the product and the ether was decanted. The product was dried under reduced pressure via lyophilization to yield a red crystalline powder.

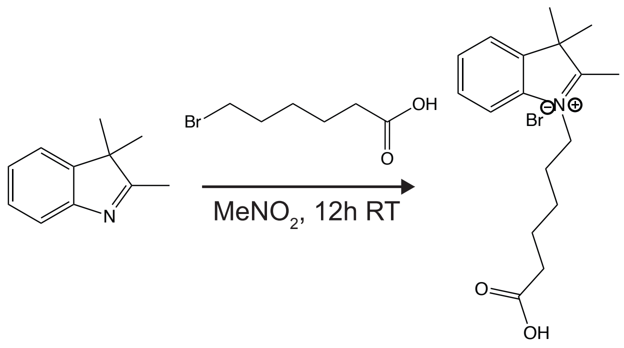


*Synthesis of carboxylated indolenine*: 6-bromohexanoic acid (1.0 mol) was added to a solution of the 2,3,3-trimethylindolenine (1.0 mol) in nitromethane ($MeNO_2$) (600 mL). The reaction was initially cooled on ice, then stirred for 24 hours at room temperature. The reaction mixture was then triturated with diethyl ether in 50 mL tubes, centrifuged to concentrate the product, and the ether was decanted. The product was then dried under vacuum on a lyophilizer. [1]

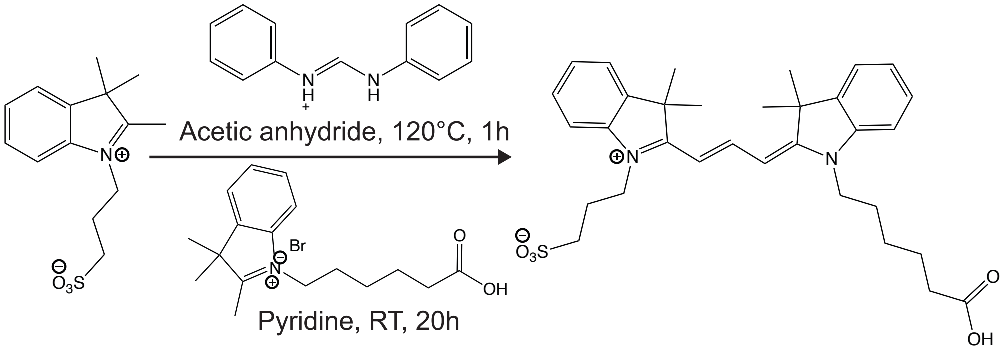


*Synthesis of Cy3*: The sulfated indolenine (50.0 mmol) and *N*,*N*-diphenylformamidine (60.0 mmol) were dissolved in acetic anhydride (150 mL) and heated at 120 °C for 30 min in an oil bath. The reaction was then cooled to room temperature and a solution of the carboxylated indolenine (70 mmol) dissolved in dry pyridine (150 mL) was added. The mixture was stirred at room temperature for 12 h and concentrated. The residue was then dissolved in chloroform (100 mL) and precipitated in diethyl ether (1 L). The residue was purified using silica gel chromatography using methanol in dichloromethane as eluent (gradient of 10-20%).

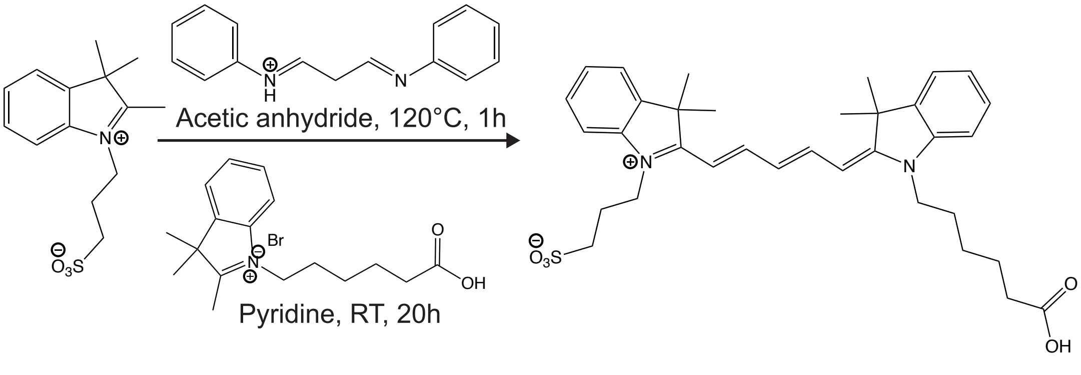

*Synthesis of Cy5*: The sulfated indolenine (50.0 mmol) and malondialdehyde bis(phenylimine) monohydrochloride (60.0 mmol) were dissolved in acetic anhydride (150 mL) and heated at 120 °C for 30 min in an oil bath. The reaction was then cooled to room temperature and a solution of the carboxylated indolenine dissolved in dry pyridine (150 mL) was added. The mixture was stirred at room temperature for 12 h and concentrated. The residue was then dissolved in chloroform (100 mL) and precipitated in diethyl ether (1 L). The residue was purified using silica gel chromatography using methanol in dichloromethane as eluent (gradient of 10-20%).

*Peptide Synthesis Procedure*
Peptides were synthesized using standard solid phase peptide synthesis protocols using either manual synthesis or an automated peptide synthesizer (CEM Liberty Blue) using standard Fmoc-protected amino acids (Chemscene) on a Rink amide resin (Supra Sciences) unless otherwise noted. All amide couplings were done using O-(6- chlorobenzotriazol-1-yl)-N,N,N′,N′-tetramethyluronium hexafluorophosphate (HCTU) unless otherwise noted. For each coupling the amino acid, HCTU, and DIPEA were added in a 4:4:6 ratio to the peptide. During peptide synthesis a ninhydrin test was performed after every addition to test for the presence of free amines. Upon a positive test, the coupling was replicated until the test was negative. A capping step was then performed with acetic anhydride (Sigma-Aldrich) in a 10:5:100 acetic anhydride:DIPEA:DMF solution twice for 5 min, and then a ninhydrin test was performed to check for complete capping of the free amines. After successful coupling, the Fmoc group was removed, washing the resin with 20% piperidine in DMF twice for 5 min. A ninhydrin test was performed to check for a positive result.

Fmoc-Lys(Mtt)-OH was coupled to the resin to synthesize PAs dye labelled with cyanine dyes. Lys(Mtt) was selectively deprotected with 3% TFA, 2% TIPS, and 95% DCM for 5 minutes after coupling the palmitic acid. Cyanine dyes (1 equivalent) were coupled to the lysine-free amine after being activated for 1 minute with HATU (0.95 equivalent) and DIPEA (1 equivalent).

All peptides were cleaved using 95% trifluoroacetic acid (TFA), 2.5% $H_2O$, 2.5% triisopropylsilane (TIPS). Peptides which contain a tryptophan were cleaved with 2.5% dithiothreitol (DTT). Peptides were typically cleaved for 2-3 hours at room temperature using approximately 2 mL of cleavage solution per mM of peptide. However, peptides containing azides were cleaved for 30 minutes to prevent degradation of the azide group. The mass was checked using electrospray ionization and if protecting groups remained the peptide was re-cleaved for 30 minutes. At the end of the cleavage the peptides were precipitated in diethyl ether. These were then centrifuged for 5 minutes at 4,000 rpm, and the supernatant was discarded. The peptide pellet was washed with diethyl ether, and centrifuged, and this was repeated three times. The peptide pellet was allowed to dry, and then dissolved in water and neutralized with ammonium hydroxide prior to purification.

*Peptide Amphiphile Purification*
All PAs were purified using high-performance liquid chromatography (HPLC) on a Phenomenex Gemini 5 μm NX-C18 110 Å LC Column (150 × 21.2 mm). For the PAs, the purification was conducted with a gradient elution starting from 95% Mobile Phase A (water containing 0.1% ammonium hydroxide and 20 mM ammonium formate) and 5% Mobile Phase B (acetonitrile containing 0.1% ammonium hydroxide), gradually increasing to 100% Mobile Phase B. For the remaining peptides, the gradient was run from 95% Mobile Phase A (water containing 0.1% trifluoroacetic acid (TFA)) and 5% Mobile Phase B (acetonitrile containing 0.1% TFA) to 100% Mobile Phase B. Each HPLC run included a two-minute equilibration phase, followed by a 10-minute gradient from 95% Mobile Phase A to 100% Mobile Phase B, and a subsequent two-

minute equilibration at 100% Mobile Phase B, before returning to the initial conditions. Following purification, all peptides were lyophilized and prepared for further use.

The PanMMP crosslinking peptide was functionalized with 2-azido acetic acid on the N-terminus. 2-azido acetic acid was synthesized by mixing bromoacetic acid (70.168 g, 505 mmol) and sodium azide (32.504 g, 500 mmol) and water (250ml), the solution was stirred overnight at RT under ambient conditions. The next day the solution was acidified to pH ~ 1 using hydrochloric acid and extracted using ethyl acetate (5 X 100 mL). The organic layers were combined and dried in vacuo to afford the 2-azidoacetic acid as a colorless liquid. 2-azidoacetic acid was stored in a -20°C freezer until needed for synthesis.

*Hydrogel Fabrication*
20 µL hydrogels were used for all studies. Note that $G_8E_3$GRGDS is "Fast-RGD", $V_4A_3E_3$GRGDS is "Slow-RGD", and $N_3$GRGDS is "PEG-RGD". The hydrogels were prepared with the following compositions:
Fast-RGD + PEG-RGD: 2.75% (wt/wt) PEG, 0.75 mM $N_3$GRGDS, 0.75 mM L-$G_8E_3$GRGDS, 0.75 mM D-$G_8E_3$, and 40% of the arms functionalized with $N_3$KGPQGIWGQKK($N_3$).
Slow-RGD + PEG-RGD: 2.75% (wt/wt) PEG, 0.75 mM L-$V_4A_3E_3$GRGDS, 0.75 mM $N_3$GRGDS, 0.75 mM D-$V_4A_3E_3$, and 40% of the arms functionalized with $N_3$KGPQGIWGQKK($N_3$).
PEG-RGD: 2.75% (wt/wt) PEG, 1.5 mM $N_3$RGD, 1.5 mM D-$V_4A_3E_3$, and 40% of the arms functionalized with $N_3$KGPQGIWGQKK($N_3$).
Fast-RGD: 2.75% (wt/wt) PEG, 1.5 mM L-$G_8E_3$GRGDS, and 40% of the arms functionalized with $N_3$KGPQGIWGQKK($N_3$).
Slow-RGD: 2.75% (wt/wt) PEG, 1.5 mM L-$V_4A_3E_3$GRGDS, and 40% of the arms functionalized with $N_3$KGPQGIWGQKK($N_3$).
Long Relaxation Gels: 2% (wt/wt) PEG, 0.75 mM L-$G_8E_3$GRGDS, 0.75 mM $N_3$GRGDS, and 95% of the arms functionalized with $N_3$KGPQGIWGQKK($N_3$).
Stiff Gels: 4% (wt/wt) PEG, 0.75 mM L-$G_8E_3$GRGDS, 0.75 mM $N_3$GRGDS, and 90% of the arms functionalized with $N_3$KGPQGIWGQKK($N_3$).

Human umbilical vein endothelial cells (hUVECs) at passage 2 were cultured in T-75 flasks using basal media (Lifeline Cell Technology, LM-0002) supplemented with ascorbic acid, hydrocortisone, fetal bovine serum (FBS), L-glutamine, recombinant human epidermal growth factor (rhEGF), heparin, and endothelial cell growth supplement (EnGS-US) (all supplements from LifeFactors, LS-1122) until reaching 80-90% confluency. The cells were then washed with PBS, trypsinized using 2 mL of 0.25% trypsin in HBSS with EDTA (Cytiva, SH30042.01), and incubated for 5 minutes. After detachment, the cells were centrifuged at 0.2 RCF for 5 minutes and counted using a hemocytometer. Following centrifugation and removal of the supernatant, the cells were resuspended to a final concentration of 2,000,000 cells/mL. 5 µL of cell suspension was then seeded into 20 µL of hydrogel to give a final cell concentration of 500,000 cells/mL. Hydrogels were made by mixing PEG-DBCO and the crosslinking peptide using copper-free click chemistry between the strained alkyne DBCO and azides. The mixture was pipetted until well-mixed, and after a 15-minute incubation, 1 mL of media was added to each well and stored in the incubator.

For confocal microscopy imaging, 12 mm round glass coverslips were placed in each well of 24-well plates. For cell viability assays, Sylgard layers were prepared in the 24-well plates to prevent unintended cell adhesion to the well bottoms. Sylgard Part A and Part B were mixed in a 10:1 ratio by volume, and the mixture was applied to the wells, ensuring complete coverage of the well bottoms. The plates were then placed in a biosafety hood and allowed to cure overnight under UV light.

*Immunocytochemistry*
The hydrogels were washed three times with PBS and then incubated in 4% formaldehyde in hUVEC media for 20 minutes. Following fixation, the hydrogels were washed three times with PBS and permeabilized with 0.25% Triton X-100 in PBS for 20 minutes. After three additional PBS washes, the hydrogels were incubated with 1% BSA in PBS for 20 minutes to block non-specific binding sites.

For vinculin staining, the hydrogels were incubated overnight at 4°C with the primary antibody, Anti-Vinculin antibody [EPR8185] (Abcam, Cat. No. ab18058) diluted 1:200 in PBS containing 1% BSA. The samples were then washed three times with PBS and incubated with the secondary antibody, Anti-rabbit IgG (H+L), F(ab')2 Fragment (Alexa Fluor® 488 Conjugate) (Cell Signaling Technology, Cat. No. 4412) diluted 1:400 in PBS with 1% BSA for 1 hour at room temperature in the dark.

After vinculin staining, the hydrogels were washed three times with PBS, and a Phalloidin/DAPI PBS staining solution (0.1% BSA, 1 µg/mL DAPI, and 1 µg/mL phalloidin-iFluor 555) was added to each well. This solution was incubated for 1 hour in the dark at room temperature. The hydrogels were then washed three times with PBS, and 1 mL of PBS was added to each well. The samples were stored at 4°C until ready for imaging.

*Quantification of Cell Spreading*
Hydrogels were imaged using a confocal microscope (LSM 800, AxioObserver, EC Plan-Neofluar 10x/0.3 objective). The 12 mm round slides with hydrogels were removed from the 24-well plates and inverted, placing the gel side down on round bottom slides. Confocal microscopy was performed in Z-stack mode. Phalloidin was excited at 560 nm, with emission recorded between 493 and 630 nm; DAPI was excited at 405 nm, with emission recorded between 410 and 501 nm; and Vinculin was excited at 488 nm, with emission recorded between 500 and 550 nm. Cell measurements within each hydrogel were analyzed using Imaris 10.2.0 software, employing the FilamentTracer module. Three biological replicates (n = 3) were imaged, with nine images per condition, to obtain average cell area, cell length, and fluorescence intensity in the hydrogels.

*Quantification of Cell Viability*
On Days 1, 3, and 7, the media from the wells was removed add 500 µL of a 10x diluted Alamar Blue stock solution in media was added to each well with hydrogels. Three wells without any cells were used for background subtraction. The gels were incubated for four hours, at which time the absorbance was measured with a spectrophotometer at wavelength of 579 nm (600 nm as reference). The cell viability percentage was assessed according to the manufacturer's protocol.

*Fluorescence depolarization*
An aqueous solution of 100 µL of annealed PA (final concentrations: [PA] = 6 mM, [KCl] = 3 mM, [NaCl] = 150 mM) was mixed with 2 µL of a THF solution containing 1,6-diphenyl-1,3,5-hexatriene (DPH; 1.4 mM). The mixture was incubated for 30 minutes at 25 °C. Subsequently, the mixture was diluted with an aqueous solution of KCl and NaCl (1900 µL, final concentrations: [KCl] = 3 mM, [NaCl] = 150 mM) and incubated for an additional 10-30 minutes at 25 °C, resulting in a final DPH concentration of 1.4 µM and PA concentration of 300 µM. Fluorescence measurements were performed using a Horiba Fluorolog-3 spectrofluorometer. DPH was excited at 336 nm, and emission was recorded at 450 nm. Both excitation and emission slit widths were set to 1 mm (8 nm bandwidth).

*Circular dichroism (CD) spectroscopy*

Each PA sample was diluted to 10 mM in buffer containing 150 mM NaCl and 3 mM KCl (high salt). CD spectra was recorded on a JASCO model J-815 spectropolarimeter using a quartz cell of 1 mm optical path length. Continuous scanning mode was used with a scanning speed of 100 nm per minute with the sensitivity set to standard mode. High Tension (HT) voltage was recorded for each sample to ensure that the measurement was not saturated. An accumulation of three measurements was used and a buffer sample was background-subtracted to obtain final spectra.

*Rheology*

All rheological studies used an AR-Ex 2000 rheometer. Rheology tests were performed on the gels using frequency sweeps at 1% strain from 0.1 to 10 Hz and stress relaxation performed for 30 minutes at constant strain of 15% using an 8 mm parallel plate. Each value represents the average of three-time tests.

*Förster resonance energy transfer (FRET)*

To make the single-color labelled nanofibers a stock solution of either Cy3-PA or Cy5-PA with a stock solution of unlabelled PA were prepared. Non-labelled PA (10 mM), Cy3-labelled PA (1 mM), and Cy5-labelled PA (1 mM) stock solutions were mixed in DI water, aliquoted, and kept at -20 °C. Simple mixing of PA stock solutions at the right ratio allowed precise control of the labeling of the fibers. In certain mole ratios, aqueous solutions of non-labeled PA were combined with aqueous solutions of one or both labels, then frozen at -80°C and lyophilized. The PA mixture was obtained and molecularly dissolved in TFA and TFA quickly removed by compressed air to create a homogenous solution and avoid degradation of the peptide. The resulting material was subsequently lyophilized, flash-frozen, and dissolved in NH4OH. Prior to measurements, the acquired powders were reconstituted in PBS. PA nanofibres were separately labelled with either PA-Cy3 (0.5%) or PA-Cy5 (0.5%). The two solutions were mixed and the FRET ratio, defined as the ratio between the fluorescence intensities of Cy5 acceptor and Cy3 donor. FRET can measure donor acceptor proximity, thus, the more the FRET ratio the closer two biomolecules are. Excitation wavelengths of 515 or 615 nm for Cy3 or Cy5, respectively, were used to measure the emission spectra. Excitation spectra for Cy3 and Cy5 were obtained at emission wavelengths of 578 and 676 nm, respectively. The built-in Peltier system was used to maintain the temperature at 25°C.

*Transverse relaxation nuclear magnetic resonance spectroscopy (T2-NMR)*

NMR spectra were acquired on a Brucker Avance Neo system at 500 MHz with a TCI cryoprobe. NMR spectra for PA-GRGDSs were recorded at 298 K in a 1:1 mixture of H2O/D2O as solvent, with a total PA-GRGDS concentration of 1.5mM.$^{1}$H T2-NMR measurements were performed using the CPMG_esgp2d pulse sequence, with a variable loop delay time ($d_2$0) of 1.0 ms.The initial data processing in TopSpin included phasing and baseline correction. Subsequently, the corresponding $^{1}$H peaks were fitted using T2 module in Dynamic Center from Bruker applying a two-state model.

T2 NMR experiments analyzed the relaxation rate of the methyl protons of the palmitic acid tail and the beta carbon of the D residue in the GRGDS sequence (observed at 0.76-0.81 and 3.10-3.11 parts per million [ppm], respectively). Fast-RGD showed the higher relaxation time than Slow-RGD or lower relaxation rate consistent with a greater degree of motion.

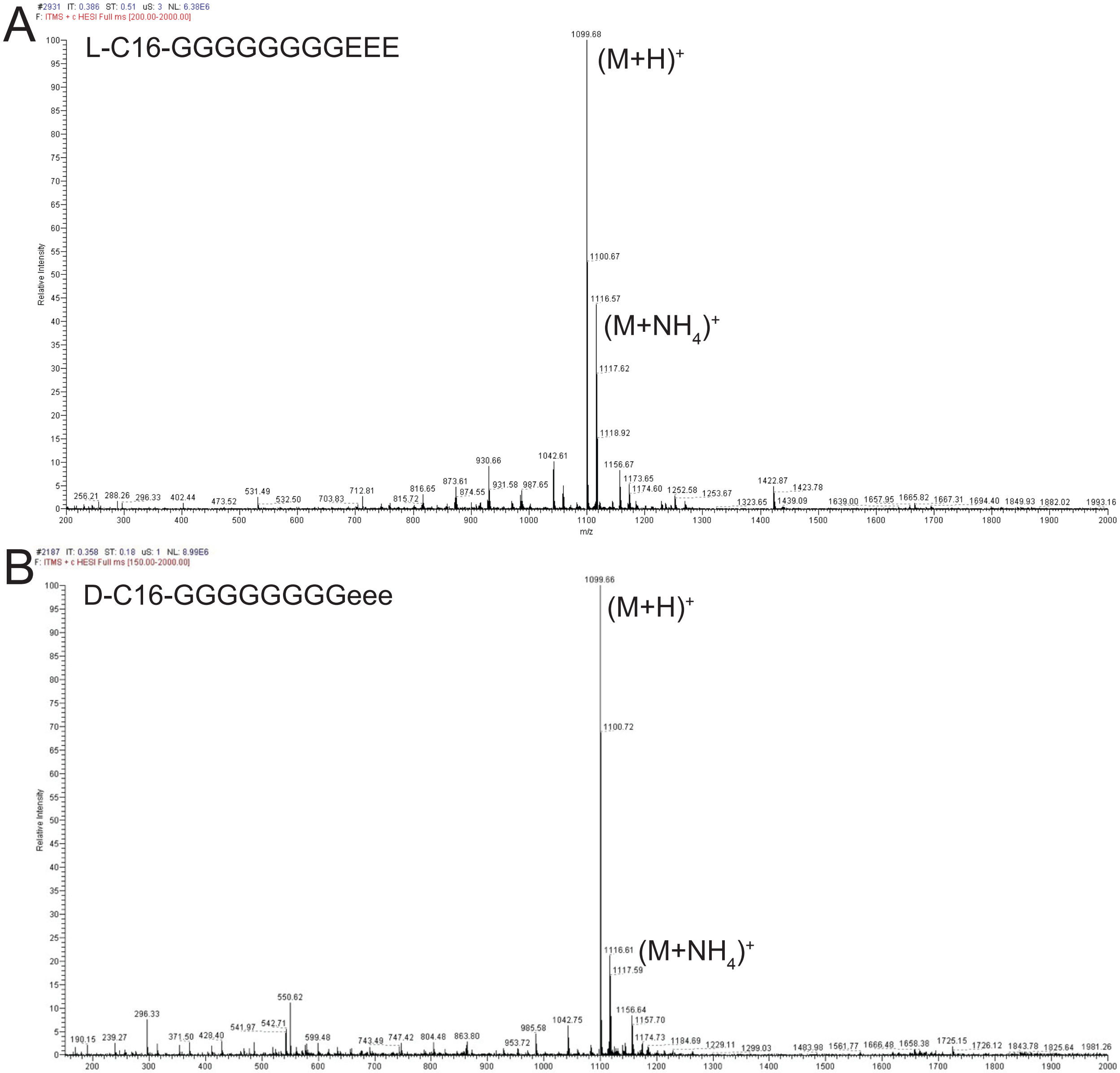
A
L-C16-GGGGGGGGEEE
(M+H)+
(M+NH4)+
1099.68
1116.57
Relative Intensity
m/z
B
D-C16-GGGGGGGGeee
(M+H)+
(M+NH4)+
1099.66
1116.61
Relative Intensity
m/z

C

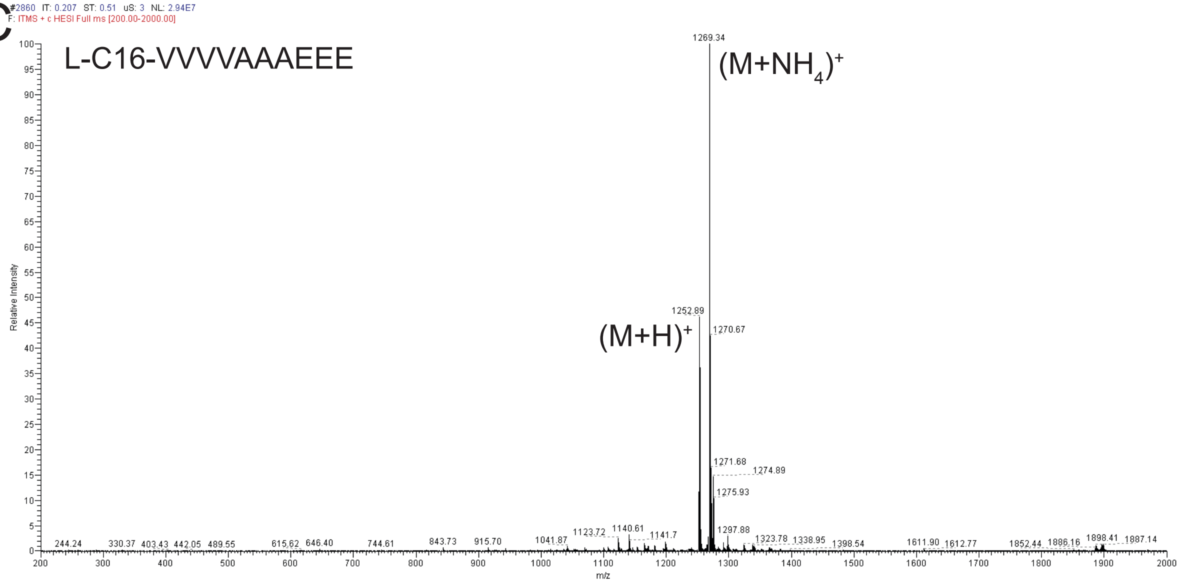
L-C16-VVVVAAAEEE
(M+NH4)+
(M+H)+
1269.34
1252.89
1270.67
Relative Intensity
m/z

D

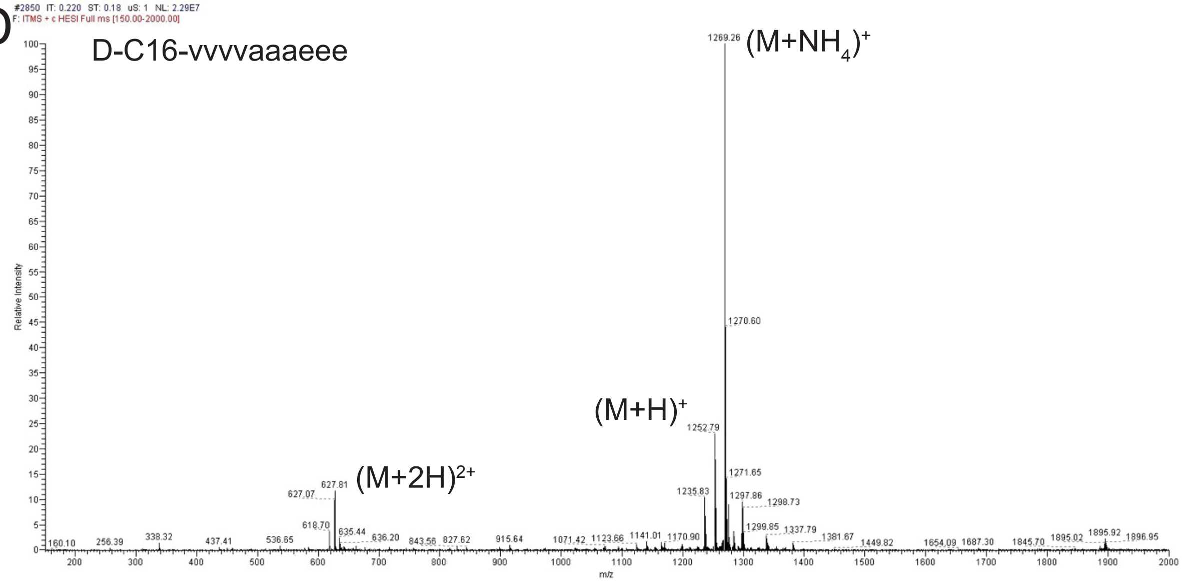
D-C16-vvvvaaaeee
(M+NH4)+
(M+H)+
(M+2H)2+
1269.26
1270.60
1252.79
627.81
Relative Intensity
m/z

E

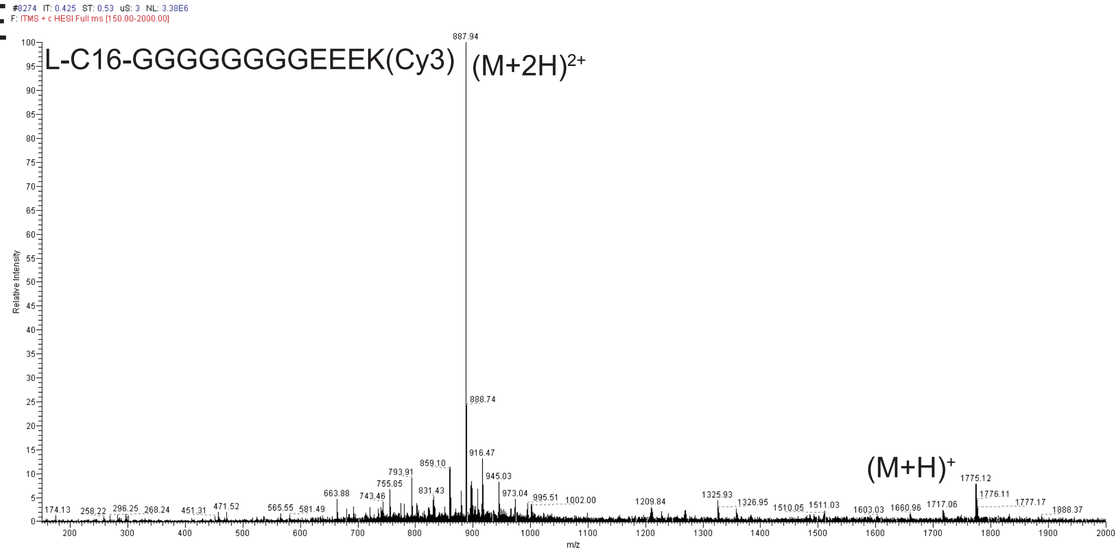
L-C16-GGGGGGGGEEEK(Cy3)
(M+2H)2+
(M+H)+
887.94
888.74
1775.12
Relative Intensity
m/z

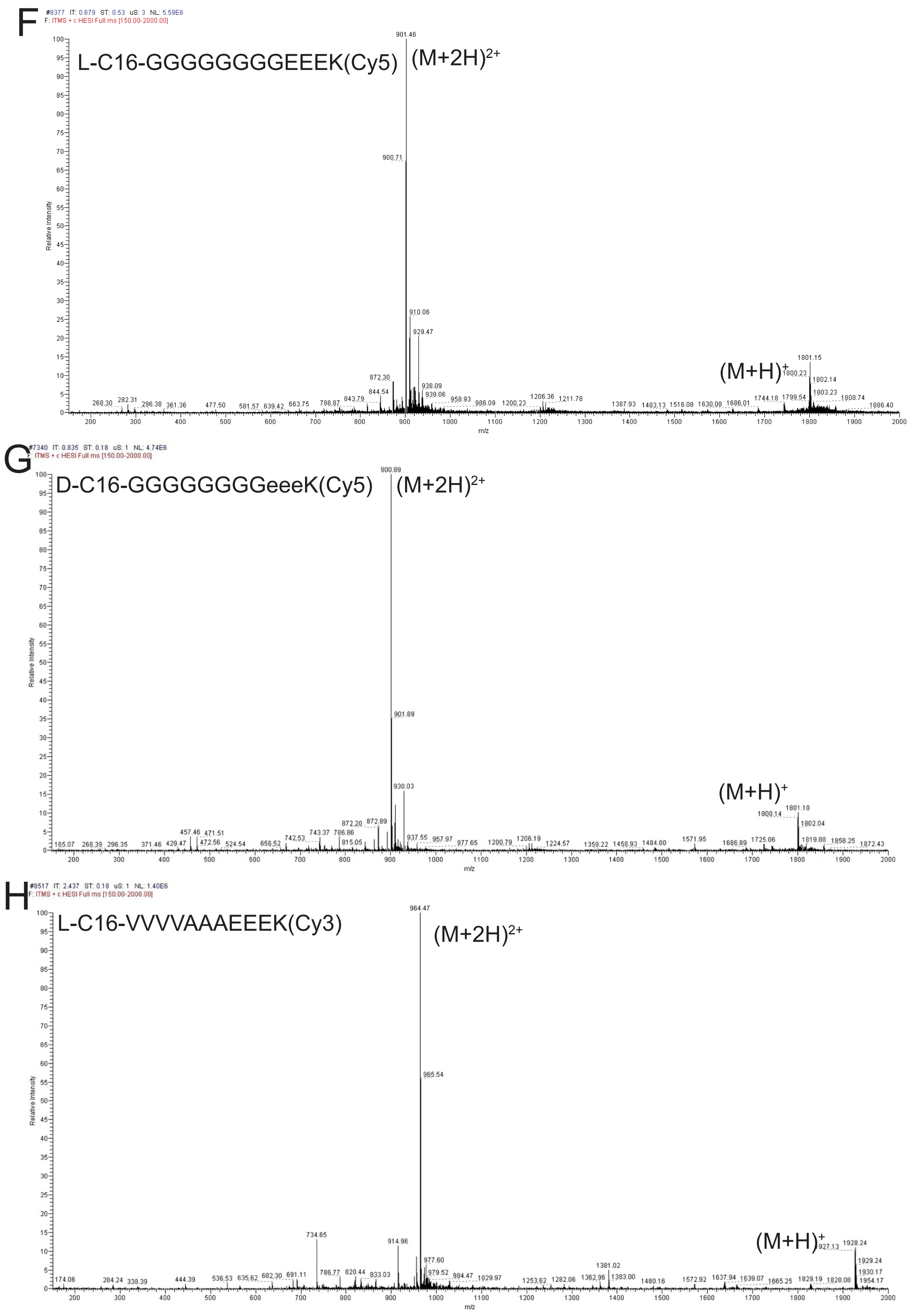
F
F: ITMS + c HESI Full ms [150.00-2000.00]
L-C16-GGGGGGGGEEEK(Cy5)
(M+2H)2+
(M+H)+
901.46
900.71
910.06
929.47
1801.15
Relative Intensity
m/z
G
F: ITMS + c HESI Full ms [150.00-2000.00]
D-C16-GGGGGGGGeeeK(Cy5)
(M+2H)2+
(M+H)+
900.89
901.89
930.03
1801.10
Relative Intensity
m/z
H
F: ITMS + c HESI Full ms [150.00-2000.00]
L-C16-VVVVAAAEEEK(Cy3)
(M+2H)2+
(M+H)+
964.47
965.54
734.65
914.96
1928.24
Relative Intensity
m/z

I

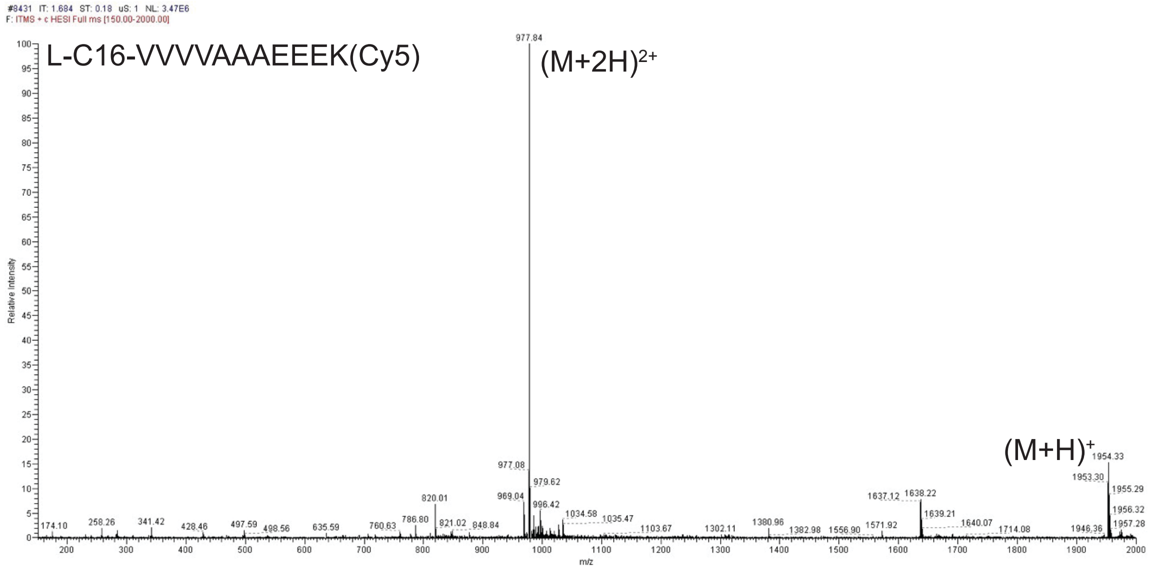
#8431 IT: 1.684 ST: 0.18 uS: 1 NL: 3.47E6
F: ITMS + c HESI Full ms [150.00-2000.00]
L-C16-VVVVAAAEEEK(Cy5)
977.84
(M+2H)$^{2+}$
(M+H)$^{+}$
1954.33
Relative Intensity
m/z

J

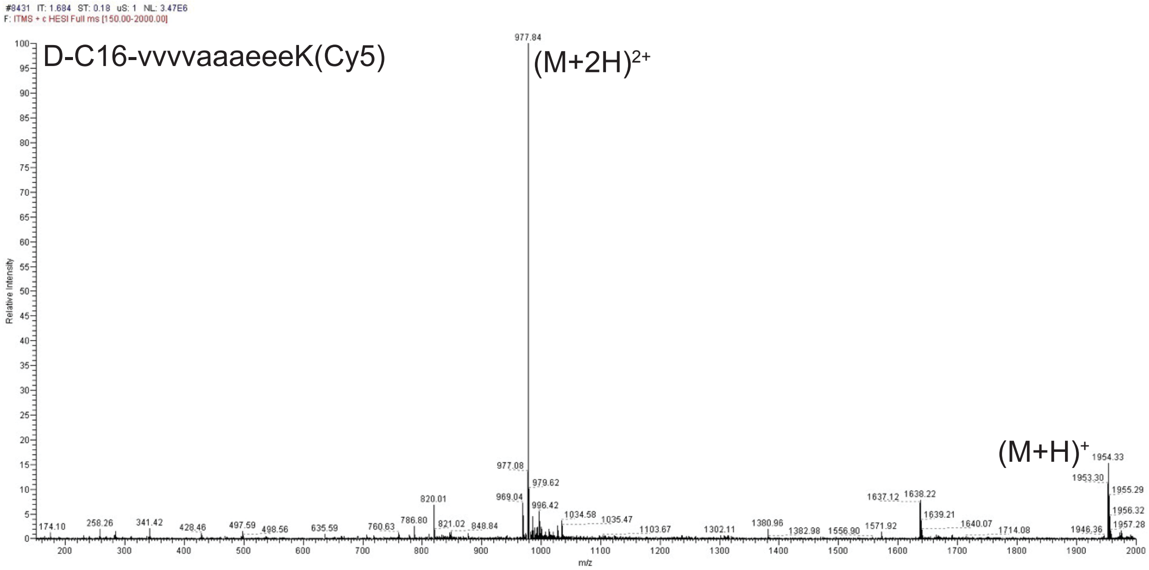
#8431 IT: 1.684 ST: 0.18 uS: 1 NL: 3.47E6
F: ITMS + c HESI Full ms [150.00-2000.00]
D-C16-vvvvaaaeeeK(Cy5)
977.84
(M+2H)$^{2+}$
(M+H)$^{+}$
1954.33
Relative Intensity
m/z

K

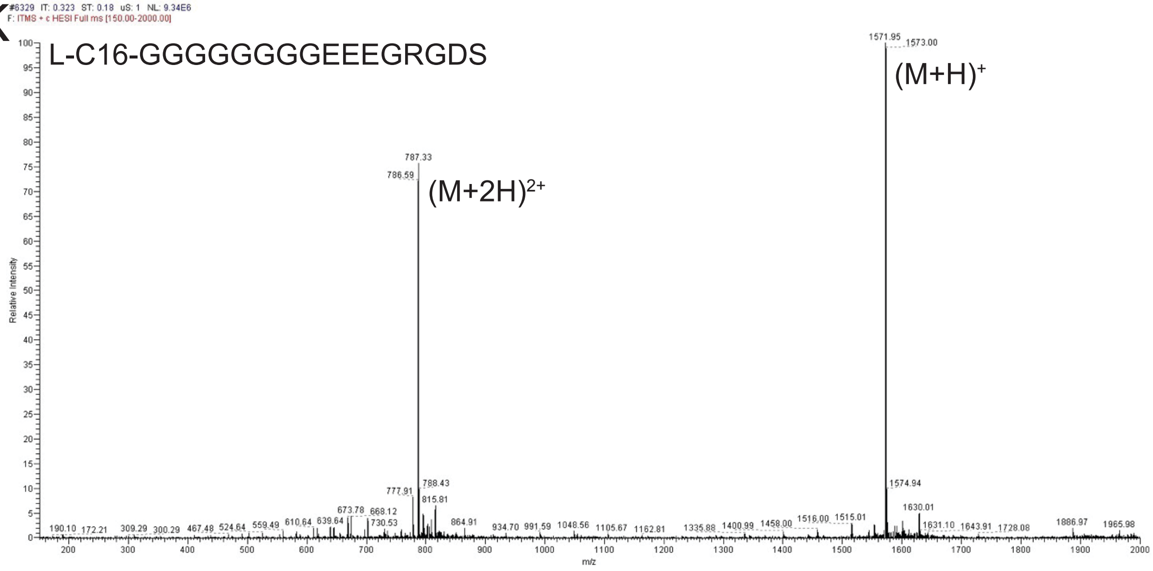
#6329 IT: 0.323 ST: 0.18 uS: 1 NL: 9.34E6
F: ITMS + c HESI Full ms [150.00-2000.00]
L-C16-GGGGGGGGEEEGRGDS
1571.95
1573.00
(M+H)$^{+}$
787.33
786.59
(M+2H)$^{2+}$
Relative Intensity
m/z

L

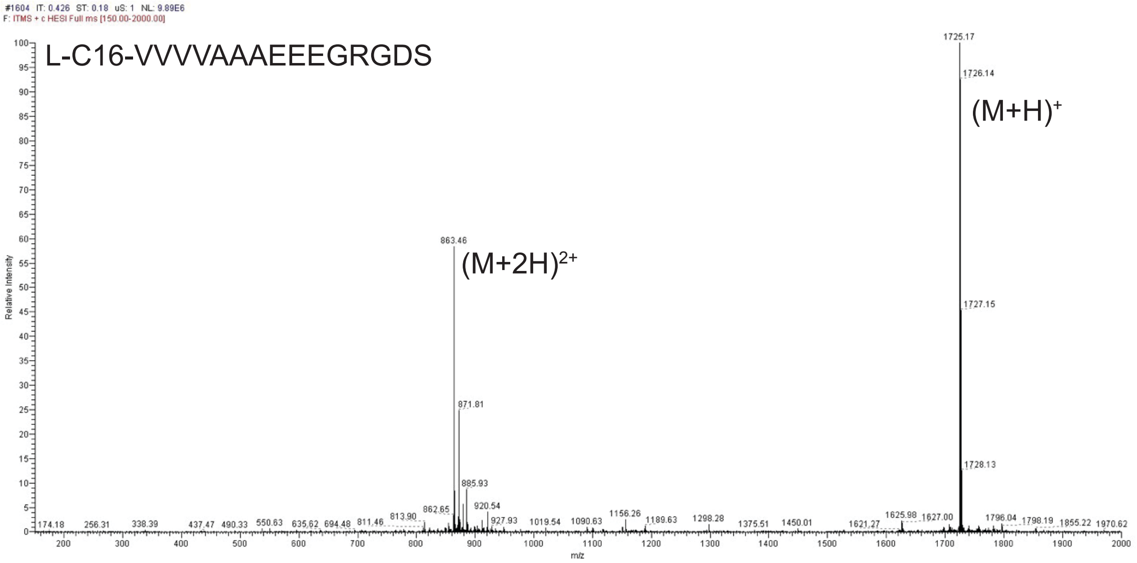
L-C16-VVVVAAAEEEGRGDS
(M+H)$^{+}$
(M+2H)$^{2+}$
1725.17
1726.14
1727.15
1728.13
863.46
871.81
885.93
920.54
Relative Intensity
m/z

M

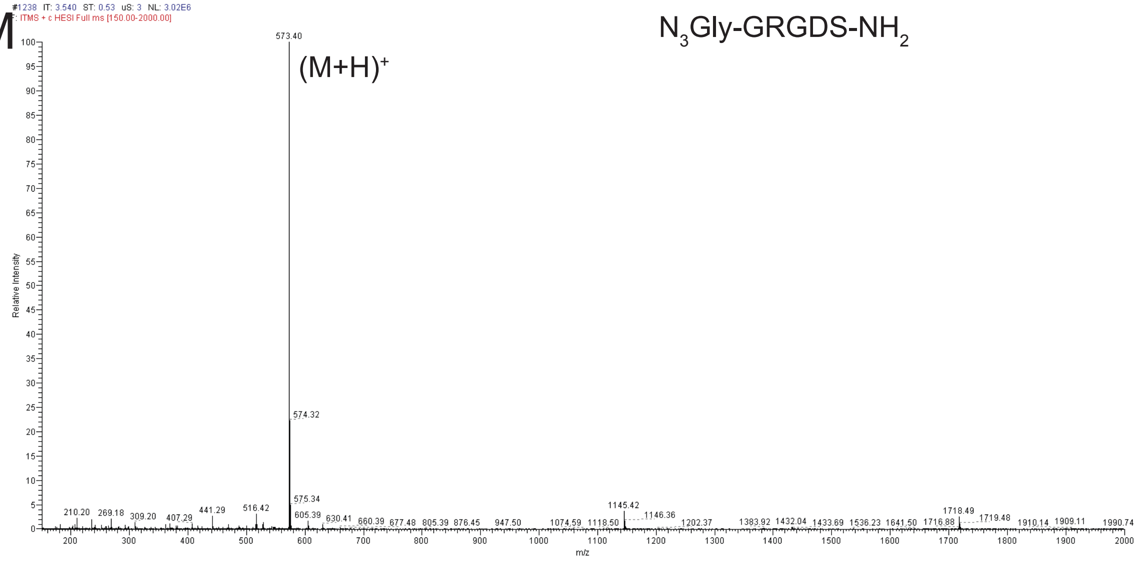
N$_3$Gly-GRGDS-NH$_2$
(M+H)$^{+}$
573.40
574.32
575.34
1145.42
Relative Intensity
m/z

N

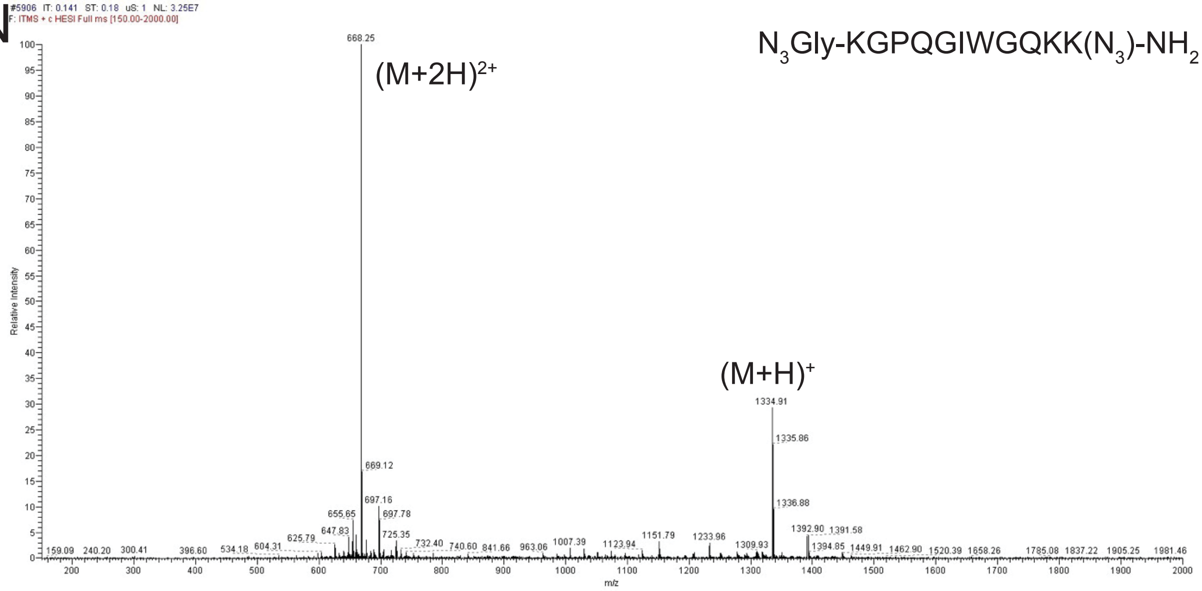
N$_3$Gly-KGPQGIWGQKK(N$_3$)-NH$_2$
(M+2H)$^{2+}$
(M+H)$^{+}$
668.25
669.12
697.16
697.78
1334.91
1335.86
1336.88
Relative Intensity
m/z

**Figure S7**. Mass spectrometry data on molecules used in these studies. A) C16-L-GGGGGGGGEEE-$NH_2$, B) C16-D-GGGGGGGGeee-$NH_2$, C) L-C16-VVVVAAAEEE, D) D-C16-vvvvaaaeee, E) L-C16GGGGGGGGEEEK(Cy3), F) L-C16GGGGGGGGEEEK(Cy5), G) D-C16GGGGGGGGeeeK(Cy5), H) L-C16-VVVVAAAEEEK(Cy3), I) L-C16-VVVVAAAEEEK(Cy5), J) D-C16-vvvvaaaeeeK(Cy5), K) L-C16-GGGGGGGGEEEGRGDS, L) L-C16-VVVVAAAEEEGRGDS, M) $N_3$-Gly-GRGDS-$NH_2$, and N) $N_3$-Gly-KGPQGIWGQKK($N_3$)-$NH_2$.

*Statistical analysis. All data was analyzed using an ANOVA followed by a pairwise comparisons using Tukey's HSD testing.*

| **Rheology** | | | | | |
|---|---|---|---|---|---|
| Condition 1 | Condition 2 | Difference | Lower 95% CI | Upper 95% CI | p-value |
| Long-relaxation Gel | Fast-RGD | -86.300000 | -216.1456 | 43.54557 | 0.2318538 |
| Slow-RGD | Fast-RGD | -77.833333 | -199.2927 | 43.62608 | 0.2564812 |
| Stiff Gel | Fast-RGD | 420.66667 | 290.8211 | 550.51224 | 0.0000158 |
| Slow-RGD | Long-relaxation Gel | 8.466667 | -112.9927 | 129.92608 | 0.9960929 |
| Stiff Gel | Long-Relaxation Gel | 506.96667 | 377.1211 | 636.81224 | 0.0000033 |

| **hUVEC Area - Day 1** | | | | | |
|---|---|---|---|---|---|
| Condition 1 | Condition 2 | Difference | Lower 95% CI | Upper 95% CI | p-value |
| Fast-RGD+PEG-RGD | Fast-RGD | 922.91933 | 546.5435 | 1299.2952 | 0 |
| PEG-RGD | Fast-RGD | 506.20889 | 129.833 | 882.58474 | 0.002349 |
| Long-relaxation Gel | Fast-RGD | 224.85467 | -151.5212 | 601.23052 | 0.536107 |
| Stiff Gel | Fast-RGD | -127.06744 | -503.4433 | 249.30841 | 0.944188 |
| Slow-RGD | Fast-RGD | 73.11467 | -303.2612 | 449.49052 | 0.996749 |
| Slow-RGD+PEG-RGD | Fast-RGD | 761.75767 | 385.3818 | 1138.1335 | 1.5E-06 |
| PEG-RGD | Fast-RGD+PEG-RGD | -416.71044 | -793.0863 | -40.33459 | 0.020895 |
| Long-relaxation Gel | Fast-RGD+PEG-RGD | -698.06467 | -1074.441 | -321.6888 | 1.04E-05 |
| Stiff Gel | Fast-RGD+PEG-RGD | -1049.9868 | -1426.363 | -673.6109 | 0 |
| Slow-RGD | Fast-RGD+PEG-RGD | -849.80467 | -1226.181 | -473.4288 | 1E-07 |
| Slow-RGD+PEG-RGD | Fast-RGD+PEG-RGD | -161.16167 | -537.5375 | 215.21418 | 0.844704 |
| Long-relaxation Gel | PEG-RGD | -281.35422 | -657.7301 | 95.02163 | 0.268932 |
| Stiff Gel | PEG-RGD | -633.27633 | -1009.652 | -256.9005 | 7.01E-05 |
| Slow-RGD | PEG-RGD | -433.09422 | -809.4701 | -56.71837 | 0.014341 |
| Slow-RGD+PEG-RGD | PEG-RGD | 255.54878 | -120.8271 | 631.92463 | 0.380604 |

| | | | | | |
|---|---|---|---|---|---|
| Stiff Gel | Long-relaxation Gel | -351.92211 | -728.298 | 24.45374 | 0.081385 |
| Slow-RGD | Long-relaxation Gel | -151.74 | -528.1159 | 224.63585 | 0.878236 |
| Slow-RGD+PEG-RGD | Long-relaxation Gel | 536.903 | 160.5271 | 913.27885 | 0.001043 |
| Slow-RGD | Stiff Gel | 200.18211 | -176.1937 | 576.55796 | 0.666143 |
| Slow-RGD+PEG-RGD | Stiff Gel | 888.82511 | 512.4493 | 1265.201 | 0 |
| Slow-RGD+PEG-RGD | Slow-RGD | 688.643 | 312.2671 | 1065.0189 | 1.38E-05 |

| **hUVEC Area - Day 7** | | | | | |
|---|---|---|---|---|---|
| Condition 1 | Condition 2 | Difference | Lower 95% CI | Upper 95% CI | p-value |
| Fast-RGD+PEG-RGD | Fast-RGD | 2407.3671 | 1316.65322 | 3498.081 | 2E-07 |
| PEG-RGD | Fast-RGD | 405.7293 | -684.984562 | 1496.4433 | 0.913577 |
| Long-relaxation Gel | Fast-RGD | 405.7293 | -684.984562 | 1496.4433 | 0.913577 |
| Slow-RGD | Fast-RGD | -606.2253 | -1696.93923 | 484.48858 | 0.619116 |
| Slow-RGD+PEG-RGD | Fast-RGD | 1099.8938 | 9.179882 | 2190.6077 | 0.046829 |
| Stiff Gel | Fast-RGD | -173.372 | -1264.0859 | 917.34192 | 0.998942 |
| PEG-RGD | Fast-RGD+PEG-RGD | -2001.638 | -3092.35168 | -910.92387 | 0.000013 |
| Long-relaxation Gel | Fast-RGD+PEG-RGD | -2001.638 | -3092.35168 | -910.92387 | 0.000013 |
| Slow-RGD | Fast-RGD+PEG-RGD | -3013.592 | -4104.30635 | -1922.8785 | 0 |
| Slow-RGD+PEG-RGD | Fast-RGD+PEG-RGD | -1307.473 | -2398.18724 | -216.75943 | 0.009342 |
| Stiff Gel | Fast-RGD+PEG-RGD | -2580.739 | -3671.45302 | -1490.0252 | 0 |
| Long-relaxation Gel | PEG-RGD | 0 | -1090.71391 | 1090.7139 | 1 |
| Slow-RGD | PEG-RGD | -1011.955 | -2102.66857 | 78.75924 | 0.085741 |
| Slow-RGD+PEG-RGD | PEG-RGD | 694.1644 | -396.549461 | 1784.8784 | 0.459578 |
| Stiff Gel | PEG-RGD | -579.1013 | -1669.81524 | 511.61257 | 0.667945 |
| Slow-RGD | Long-relaxation Gel | -1011.955 | -2102.66857 | 78.75924 | 0.085741 |
| Slow-RGD+PEG-RGD | Long-relaxation Gel | 694.1644 | -396.549461 | 1784.8784 | 0.459578 |
| Stiff Gel | Long-relaxation Gel | -579.1013 | -1669.81524 | 511.61257 | 0.667945 |
| Slow-RGD+PEG-RGD | Slow-RGD | 1706.1191 | 615.405206 | 2796.833 | 0.00025 |
| Stiff Gel | Slow-RGD | 432.8533 | -657.860572 | 1523.5672 | 0.88597 |
| Stiff Gel | Slow-RGD+PEG-RGD | -1273.266 | -2363.97968 | -182.55187 | 0.012378 |

| **Actin Intensity - Day 1** | | | | | |
|---|---|---|---|---|---|
| Condition 1 | Condition 2 | Difference | Lower 95% CI | Upper 95% CI | p-value |
| Fast-RGD+PEG-RGD | Fast-RGD | 248526.45 | 22853.44 | 474199.47 | 0.021968 |
| PEG-RGD | Fast-RGD | 268265.03 | 42592.02 | 493938.05 | 0.010225 |
| Long-relaxation Gel | Fast-RGD | -96819.78 | -322492.8 | 128853.23 | 0.843511 |
| Stiff Gel | Fast-RGD | -181086.3 | -406759.3 | 44586.74 | 0.196349 |
| Slow-RGD | Fast-RGD | -79995.54 | -305668.6 | 145677.47 | 0.930248 |
| Slow-RGD+PEG-RGD | Fast-RGD | 237539.18 | 11866.16 | 463212.2 | 0.032918 |
| PEG-RGD | Fast-RGD+PEG-RGD | 19738.58 | -205934.4 | 245411.6 | 0.999967 |
| Long-relaxation Gel | Fast-RGD+PEG-RGD | -345346.2 | -571019.3 | -119673.2 | 0.000358 |
| Stiff Gel | Fast-RGD+PEG-RGD | -429612.7 | -655285.8 | -203939.7 | 0.000006 |
| Slow-RGD | Fast-RGD+PEG-RGD | -328522 | -554195 | -102849 | 0.000774 |
| Slow-RGD+PEG-RGD | Fast-RGD+PEG-RGD | -10987.27 | -236660.3 | 214685.75 | 0.999999 |
| Long-relaxation Gel | PEG-RGD | -365084.8 | -590757.8 | -139411.8 | 0.000141 |
| Stiff Gel | PEG-RGD | -449351.3 | -675024.3 | -223678.3 | 2.2E-06 |
| Slow-RGD | PEG-RGD | -348260.6 | -573933.6 | -122587.6 | 0.000312 |
| Slow-RGD+PEG-RGD | PEG-RGD | -30725.85 | -256398.9 | 194947.17 | 0.999564 |
| Stiff Gel | Long-relaxation Gel | -84266.49 | -309939.5 | 141406.52 | 0.912129 |
| Slow-RGD | Long-relaxation Gel | 16824.24 | -208848.8 | 242497.26 | 0.999987 |
| Slow-RGD+PEG-RGD | Long-relaxation Gel | 334358.97 | 108685.95 | 560031.98 | 0.000594 |
| Slow-RGD | Stiff Gel | 101090.73 | -124582.3 | 326763.75 | 0.815181 |
| Slow-RGD+PEG-RGD | Stiff Gel | 418625.46 | 192952.44 | 644298.47 | 1.04E-05 |
| Slow-RGD+PEG-RGD | Slow-RGD | 317534.72 | 91861.71 | 543207.74 | 0.001269 |

| **Vinculin Intensity - Day 1** | | | | | |
|---|---|---|---|---|---|
| Condition 1 | Condition 2 | Difference | Lower 95% CI | Upper 95% CI | p-value |
| Fast-RGD+PEG-RGD | Fast-RGD | 142956.53 | -54992.51 | 340905.583 | 0.307639 |
| PEG-RGD | Fast-RGD | -126182.5 | -324131.5 | 71766.587 | 0.457623 |
| Long-relaxation Gel | Fast-RGD | -203417.6 | -401366.6 | -5468.536 | 0.040243 |
| Stiff Gel | Fast-RGD | -303057.6 | -501006.6 | -105108.53 | 0.000355 |
| Slow-RGD | Fast-RGD | -161721.6 | -359670.7 | 36227.441 | 0.179702 |
| Slow-RGD+PEG-RGD | Fast-RGD | 109138.1 | -88810.95 | 307087.149 | 0.627956 |

| | | | | | |
|---|---|---|---|---|---|
| Fast-RGD+PEG-RGD | Fast-RGD+PEG-RGD | -269139 | -467088 | -71189.948 | 0.002034 |
| Long-relaxation Gel | Fast-RGD+PEG-RGD | -346374.1 | -544323.2 | -148425.07 | 3.35E-05 |
| Stiff Gel | Fast-RGD+PEG-RGD | -446014.1 | -643963.2 | -248065.07 | 1E-07 |
| Slow-RGD | Fast-RGD+PEG-RGD | -304678.1 | -502627.2 | -106729.09 | 0.000326 |
| Slow-RGD+PEG-RGD | Fast-RGD+PEG-RGD | -33818.43 | -231767.5 | 164130.614 | 0.99841 |
| Long-relaxation Gel | PEG-RGD | -77235.12 | -275184.2 | 120713.926 | 0.893835 |
| Stiff Gel | PEG-RGD | -176875.1 | -374824.2 | 21073.932 | 0.109068 |
| Slow-RGD | PEG-RGD | -35539.15 | -233488.2 | 162409.902 | 0.997901 |
| Slow-RGD+PEG-RGD | PEG-RGD | 235320.56 | 37371.51 | 433269.611 | 0.010219 |
| Stiff Gel | Long-relaxation Gel | -99639.99 | -297589 | 98309.054 | 0.720218 |
| Slow-RGD | Long-relaxation Gel | 41695.98 | -156253.1 | 239645.025 | 0.994936 |
| Slow-RGD+PEG-RGD | Long-relaxation Gel | 312555.68 | 114606.64 | 510504.733 | 0.000214 |
| Slow-RGD | Stiff Gel | 141335.97 | -56613.08 | 339285.019 | 0.320812 |
| Slow-RGD+PEG-RGD | Stiff Gel | 412195.68 | 214246.63 | 610144.727 | 8E-07 |
| Slow-RGD+PEG-RGD | Slow-RGD | 270859.71 | 72910.66 | 468808.756 | 0.001867 |
